# Distinct functional roles and connectivity rules for lower and higher order intracortical and pulvinar thalamocortical pathways in mouse visual cortex

**DOI:** 10.1101/2023.05.07.539734

**Authors:** Xu Han, Vincent Bonin

**Affiliations:** 1Neuro-Electronics Research Flanders, Kapeldreef 75, 3001 Leuven, Belgium; KU Leuven, Department of Biology, 3000 Leuven, Belgium; VIB-KU Leuven Center for Brain & Disease Research, 3000 Leuven, Belgium (current address); VIB, 3001 Leuven, Belgium; imec, 3001 Leuven, Belgium; KU Leuven, Leuven Brain Institute, 3000 Leuven, Belgium

## Abstract

Functional specialization of cortical areas is a fundamental feature of brain organization and is critical for perception and behavior. Such an organization must depend on specialized connectivity between areas, yet the underlying wiring rules remain unclear. We characterized intracortical and thalamocortical pathways in the mouse visual cortex using neural tracing and functional imaging. We uncovered multiple structural-functional connectivity rules underlying the functional specialization of higher visual cortical areas (HVAs). Individual higher visual areas integrate specific cortical and thalamic inputs with distinct functional biases. Higher order Layer 2/3 and thalamocortical pathways show higher target specificity than feedforward intracortical pathways and might impart specific functional preferences to the recipient HVAs. In contrast, higher order Layer 5 pathways lacking specificity may contribute to the tuning diversity in the recipient HVAs. Altogether, this study reveals fundamental organization rules of long-range interareal connectivity that underlie the parallel modular organization of the visual cortex.

**HIGHLIGHTS:**

- HVAs AL, PM and A receive diverse and specific inputs from V1, HVA and LP pathways
- Density of intracortical inputs correlates with similarity of tuning between visual areas
- Tuning of HVA inputs correlates with HVA’s preferences and functional heterogeneity
- HVA output pathways differ in tuning homogeneity and target specificity

## INTRODUCTION

Functional specialization of cortical areas is a hallmark of the mammalian neocortex. The visual cortex, in particular, features dozens of specialized cortical areas dedicated to visual processing (Felleman and Van Essen, 1991). These higher-order visual areas (HVAs) exhibit specialized neuronal response properties and neural representations crucial for visual perception and behavior (Maunsell and Newsome, 1987; Orban, 2008). HVAs of specific visual functions are organized into segregated visual processing streams (DeYoe and Van Essen, 1988; Goodale and Milner, 1992; Mishkin et al., 1983). These interconnected networks of visual areas process distinct visual scene attributes (Hebart and Hesselmann, 2012; Priebe et al., 2003; Tanaka, 1996) and perform the visual computations necessary for perceptual, cognitive and motor behavior.

While the response properties of HVAs and associated neuronal pathways have been characterized in increasing detail, the precise visual information flows that underlie the generation of specialized representations in HVAs remain to be elucidated. Initiated in the primary visual cortex (V1), the visual streams are thought to emerge via specific innervation of thalamocortical (Lennie, 1980) and intracortical visual pathways (Maunsell and Newsome, 1987; Nassi and Callaway, 2009). However, HVA activity does not depend exclusively on feedforward pathways (Nassi et al., 2006; Tohmi et al., 2014) receiving ample inputs from higher order, HVA-to-HVA, intracortical pathways and pulvinar thalamocortical pathways (Bridge et al., 2016; Van Essen, 2005; Felleman and Van Essen, 1991; Padberg and Krubitzer, 2006). In addition, pulvinar neurons exhibit a topographic relationship with HVAs and can drive the activity of HVA neurons (Kaas and Lyon, 2007; Yao et al., 2023).

Imaging studies in mice have begun to unravel the large-scale cellular connectivity patterns underlying representational biases in HVAs. As a population, HVA neurons form distributed visual representations and visual spatiotemporal information channels (Andermann et al., 2011; Han et al., 2022; Marshel et al., 2011; Murakami et al., 2017) which can be used to probe functionally specific neural circuitry. Intracortical pathways from V1 to HVAs were found to show in responses to spatiotemporal stimuli target-specific functional tuning properties biased in the direction of the tuning properties of HVAs (Blot et al., 2021; Glickfeld et al., 2013; Kim et al., 2018; Murgas et al., 2020). However, the number of presynaptic intracortical projection neurons in each source area can vary up to three orders of magnitude (Gămănuţ et al., 2018; Wang et al., 2012), suggesting different pathways have distinct contributions to interareal communication. At a finer resolution, interareal communication pathways are composed of multiple ‘lanes’ that are mediated by distinct subtypes of intracortical projection neurons with distinct cortical laminar distributions and axonal projection profiles (Han et al., 2018; Harris et al., 2019). Whether these parallel pathways mediated by distinct intracortical subpopulations are organized by similar rules (e.g., functionally target-specific) and play similar functional roles remains unclear.

Silencing of V1 inputs only mildly impairs mouse HVA visual representations (Tohmi et al., 2014), suggesting the necessity of additional input pathways. The mouse lateral posterior thalamic nucleus (LP), the analogous structure of the primate pulvinar, comprises multiple clusters of neurons that project onto distinct HVAs (Bennett et al., 2019; Juavinett et al., 2020). In addition, different LP subregions show different responses to visual spatiotemporal frequencies and visual motion (Beltramo and Scanziani, 2019; Bennett et al., 2019; Blot et al., 2021), suggesting parallel pathways conveying different streams of spatiotemporal information from LP to different HVAs. However, the functional organization of these higher order thalamocortical pathways remains to be systematically characterized, and a complete account of the functional tuning of the diverse pathways converging onto mouse HVAs is still lacking (Gămănuţ and Shimaoka, 2021).

In this study, we combined genetic circuit labeling with widefield microscopy and cellular imaging to characterize the joint anatomical and functional organization of input and output pathways to and from three HVAs, probing the visual tuning of lower and higher order L2/3 and L5 intracortical pathways to areas AL, PM, and A, and of LP thalamocortical pathways to HVAs. At the mesoscopic level, we found a correlation between the areas’ response properties and the density of intracortical long-range inputs. At the cellular level, however, we found diverse intracortical connectivity patterns. While AL and PM integrate diverse visual inputs from diverse sources, area A receives much more homogeneous and stereotyped visual information. Furthermore, pathways to HVAs vary widely in their degree of target specificity, with higher order L2/3 pathways and LP pathways showing the highest degree of target specificity and higher order L5 pathways showing the least specificity. The properties of the pathways suggest higher order HVA and LP pathways obey distinct functional roles and play prominent albeit specific roles in shaping the functional diversity and specificity of HVAs.

## RESULTS

### Heterogeneous spatiotemporal tuning underlies specialized HVA representations

A diversity of neuronal pathways and functional response properties are suggested to underlie specialized representations and processing streams in the visual cortex (Gămănuţ and Shimaoka, 2021; Nassi and Callaway, 2009). In the mouse, higher visual cortical areas (HVAs) show specific spatiotemporal tuning properties (Andermann et al., 2011; Han et al., 2022; Marshel et al., 2011; Murakami et al., 2017) and are innervated by specific pathways (Blot et al., 2021; Glickfeld et al., 2013; Han et al., 2018; Kim et al., 2018). To shed light on the joint functional and anatomical pathway organization giving rise to specialization, we characterized, using *in vivo* functional cellular imaging and neural circuit tracing, the spatiotemporal receptive field properties of intracortical and thalamocortical neurons projecting to and from specific HVAs (Figure 1A).

**Figure 1.**
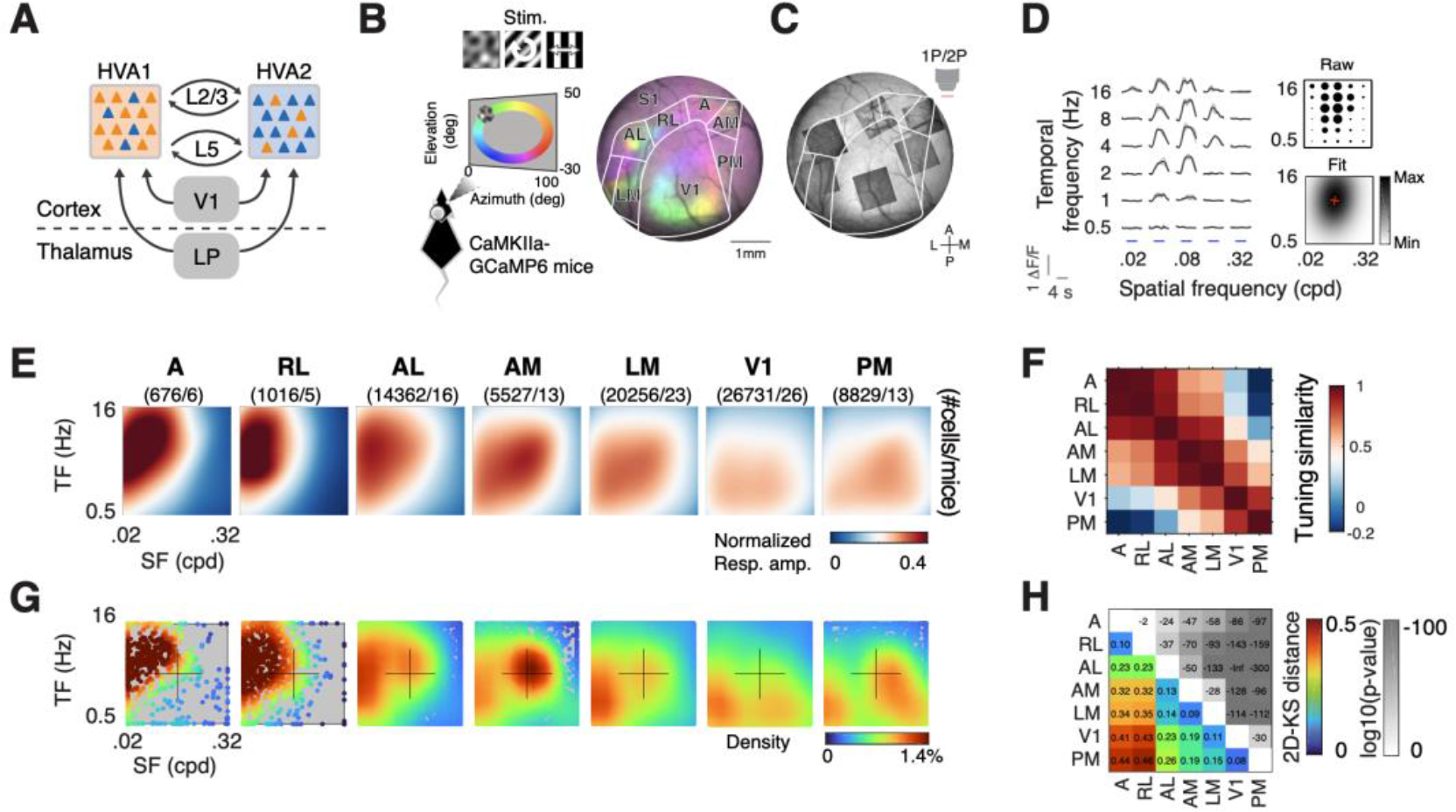
Heterogeneous spatiotemporal tuning underlies specialized HVA representations. (A) Schematic showing the intracortical and thalamocortical communication pathways between V1, lateral posterior thalamic nucleus (LP), and higher visual areas (HVAs). (B) Schematic showing the visual stimulation paradigm and an example retinotopic map over the surface of the left visual cortex of a GCaMP6- transgenic mouse (CaMKII-tTA x TRE-GcaMP6s), with areal delineation (white contours). The cortex imaged through a chronic cranial window is color-coded by the location of the visual field which generated the highest neural responses. (C) Example two-photon fields-of-view over different visual cortical areas, superimposed on a one-photon image of the cortical surface. (D) Spatiotemporal frequency tuning of an example cortical neuron. Traces are trial- averaged calcium fluorescence changes in response to visual stimuli with distinct combinations of spatial and temporal frequencies. The largest mean response amplitudes (Raw, top right) evoked by the visual stimulus sets were fitted to a 2D Gaussian model (bottom right). Red cross: The neuron’s preferred (peak) spatiotemporal frequency. (E) Distinct averaged spatiotemporal tuning surfaces of neurons across areas. The tuning surfaces were the averages of different subsampling of datasets across animals with hierarchical bootstrapping. The numbers of cells and mice are shown at the top. (F) Tuning similarity across visual cortical areas. Tuning similarity is calculated as Pearson’s correlation coefficients between average tuning maps. (G) Distinct distributions of peak spatial (SF) and temporal frequencies (TF) of neurons in different areas. (H) Pairwise distances of the 2D distributions of peak spatiotemporal frequencies. The lower left triangle shows two-dimensional Kolmogorov–Smirnov (2D-KS) distances. P-values are shown in the upper right triangle. See also Figure S1.

To get a comprehensive account of functional specialization in mouse visual areas, we characterized in TRE-GcaMP6s x CaMKII-tTA mice (N = 28 mice) (Wekselblatt et al., 2016) the spatiotemporal tuning preferences of >70,000 L2/3 excitatory neurons distributed across eight retinotopic visual areas. Previous studies of HVAs have shown tuning diversity both within and across visual areas (Andermann et al., 2011; Han et al., 2022; Marshel et al., 2011; Murakami et al., 2017; de Vries et al., 2019). These differences seem partially reflected in the input pathways (Blot et al., 2021; Glickfeld et al., 2013; Kim et al., 2018). To capture that diversity, we recorded in HVAs responses to three distinct types of visual stimuli sampling 30 combinations of spatial and temporal frequencies, two degrees of spatial anisotropy, and four spatial orientations and directions of motion direction (Figure 1B, S1A-D) (see Star Methods). Visual areas were delineated using widefield calcium imaging (Figure 1B). V1, lateromedial (LM), anterolateral (AL), rostrolateral (RL), anterior (A), anteromedial (AM), and posteromedial (PM) (Wang and Burkhalter, 2007) were targeted with 2-photon (2P) recordings (Figure 1C). Cellular calcium responses were measured in the superficial cortical layer (L2/3, 100-300 um depths), and reliable visually tuned neurons were selected using correlational analysis (Figure S1F, cross-trial correlation > 0.3, see Star Methods).

The different types of stimuli yielded consistent tuning but evoked different response magnitudes (Figure S1A-D). For each neuron, we thus selected the measurements from the stimulus-response set with the highest peak response magnitudes (Figure S1C, thick lines). Then, using 2-dimensional Gaussian function fits applied onto the average calcium responses over stimulation epochs (Figure 1D, S1B-C), we computed the neurons’ mean average response amplitudes, the preferred (peak) spatial and temporal frequencies, and the preferred (peak) speed (i.e., the spatial and temporal frequencies, and temporal-to-spatial frequency ratios eliciting the largest responses).

We first examined the neurons’ average response to the stimuli for each area. We observed widely distinct spatiotemporal tuning biases across visual areas (Figure 1E), consistent with recordings in a different mouse line with GCaMP6 targeted to a subset of excitatory neurons (Han et al., 2022). While V1 shows the strongest activation for stimuli of low temporal frequencies, LM shows a peak at mid-range spatial and temporal frequencies and weak responses at high spatial frequencies, which is not seen in V1. In contrast, anterior visual areas (AL, RL, A and AM) are preferentially activated by stimuli of low spatial and high temporal frequencies (0.02-0.08 cpd, 4- 16 Hz), whereas the posterior area PM shows a preference for low-temporal, high-spatial frequency stimuli (0.04-0.16 cpd, 1-4Hz). As visible in the pairwise correlation matrix of the neurons’ average tuning maps, distinct tuning biases are observed in anterior and posterior HVAs (Figure 1F), suggesting HVAs may receive distinct visual information from different pathways.

We next examined the neurons’ preferred spatiotemporal frequencies and observed a high diversity of tuning preferences in HVAs. Most areas differ in their distributions of peak spatial and temporal frequency, and peak speed (Figure S1G). These differences are robust to the correlation thresholds used to select visually tuned neurons (Figure S1F). In many areas, peak 2-dimensional spatiotemporal frequencies distributions are separable with distinct modes at distinct spatiotemporal frequencies (Figure 1G). As observed for the average tuning, anterior and posterior HVAs show highly distinct spatiotemporal distributions (Figure 1 G, H). Within visual areas, neurons also show dispersed tuning preferences, as observed in most areas (V1, LM AL, AM, and PM). Although responses in A and RL appear less dispersed, the sample sizes for these areas were smaller. This high diversity suggests HVAs may receive diverse visual information from afferent pathways.

### Similarly tuned visual areas provide highest density of intracortical inputs to HVAs

The tuning properties of HVA neurons suggest HVAs integrate specific and diverse visual inputs from other visual areas. To characterize multiway intracortical input pathways innervating HVAs, we performed injections of retrograde AAV (retroAAV-CAG-tdTomato) (Tervo et al., 2016) in GCaMP6 mice (Figure 2A-B). We targeted AL, PM and A, which display distinct functional properties and receive distinct pathway innervation (D’Souza et al., 2022; Yao et al., 2022) and, therefore, should integrate distinct sets of inputs (Blot et al., 2021; Glickfeld et al., 2013; Kim et al., 2018). To characterize the magnitude of cortical inputs received from other visual areas, we captured in vivo widefield 1-photon (1P) fluorescence images of the cortical surface and examined tdTomato labeling patterns (Figure 2B-C). To estimate the magnitude of inputs from individual visual areas, we calculated the total fluorescence intensity over each non-injected area and normalized the result to the highest values across non-injected areas (Figure 2D, normalized total fluorescence). 2P image stacks captured in vivo confirmed that this connectivity measure correlates with the density of retrogradely-labeled somas (Figure S2).

**Figure 2.**
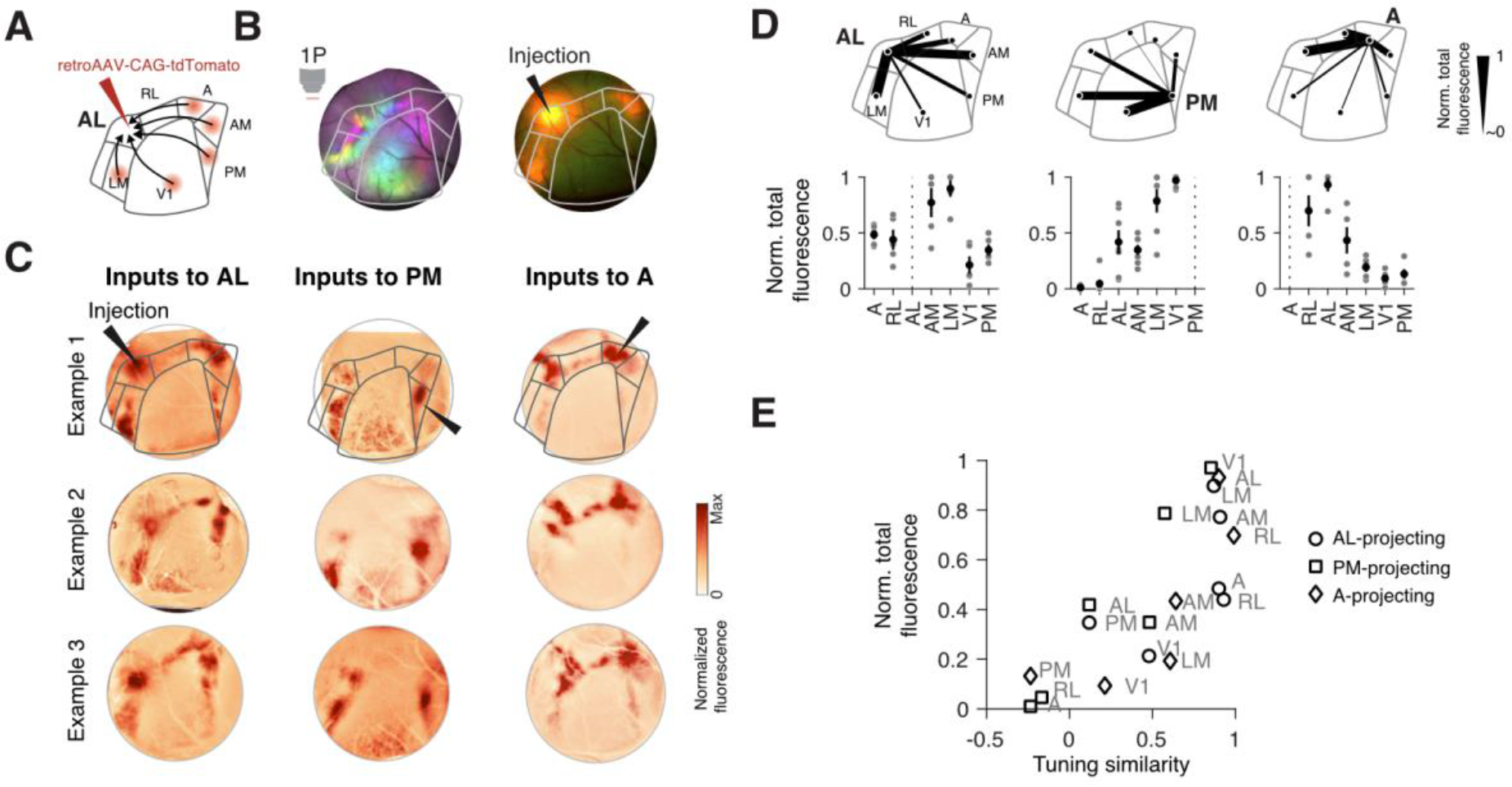
HVA inputs originate preferentially from similarly tuned visual cortical areas. (A) Schematic of retrograde tracing of intracortical input pathway. (B) Widefield images of the cortical surface showing the retinotopic map (left) and tdTomato expression 10 days after the injection of the retrograde virus at area AL (right). (C) Example cortical input maps resulting from injections of retrograde viruses in different areas (columns) in different mice (rows). Gray contours: area delineation. (D) Summary of the anatomical intensity of cortical input pathways to areas AL, PM, and A. N=5,8,5 mice, respectively (gray markers). Black markers: mean ± SEM (E) Scatter plot showing the correlation between the strength of retrograde intracortical labeling and the similarity of visual tuning amongst connected HVAs. The source areas are shown in the text. The target areas are represented as different types of markers. See also Figure S2.

These experiments revealed area-specific, functionally related intracortical projection patterns to HVAs (Figure 2D-E). Consistent with previous anatomical work (D’Souza et al., 2022; Yao et al., 2022), HVAs receive inputs from distinct subsets of visual areas (Figure 2C-D). While AL and PM receive inputs from a broad set of visual cortical areas, including V1, area A’s intracortical innervation is more specific and restricted mainly to anterior HVAs. Furthermore, the strength of innervation strongly depends on presynaptic area anatomical and functional identity. While AL and A receive their strongest inputs from anterior HVAs, area PM receives its strongest inputs from posterior HVAs. Interestingly, the density of intracortical projection pathways and the similarity of functional tuning are correlated (Figure 2E), with AL, PM, and A receiving their strongest innervation from similarly tuned visual areas. Together, these data indicate HVAs receive specific intracortical inputs preferentially integrating information from similarly tuned visual areas.

### Distinct connectivity rules amongst L2/3 intracortical neurons projecting to HVAs

Having observed a general link between functional tuning and the density of intracortical projection pathways, we next characterized the receptive field properties of intracortical neurons projecting to HVAs. Previous studies suggested specialized HVA representations result from the convergence of inputs from lower visual areas reporting in V1 and LM tuning biased towards the tuning of target HVAs (Glickfeld et al., 2013; Kim et al., 2018). To assess the distinct contributions of distinct pathways, including pathways from lower and higher visual areas, we characterized using 2P imaging in GCaMP6 mice the spatiotemporal visual tuning of V1 and HVA neurons projecting to AL, PM or A (Figure 3A-C, N=9, 13, 5 mice, respectively). We compared the tuning of tdTomato+ retrogradely-labeled neurons (tdT+) and non-labeled neurons (tdT–) in the source area to the tuning of the populations in the target area. We asked whether the neurons show specialization, conveying diverse and specific visual information to HVAs. We considered two types of specificity: source specificity, whereby neurons from different source areas convey distinct visual information to a specific target, and target specificity, whereby projection neurons from the source area anatomically and functionally diverge, showing tuning like that of their targets.

**Figure 3.**
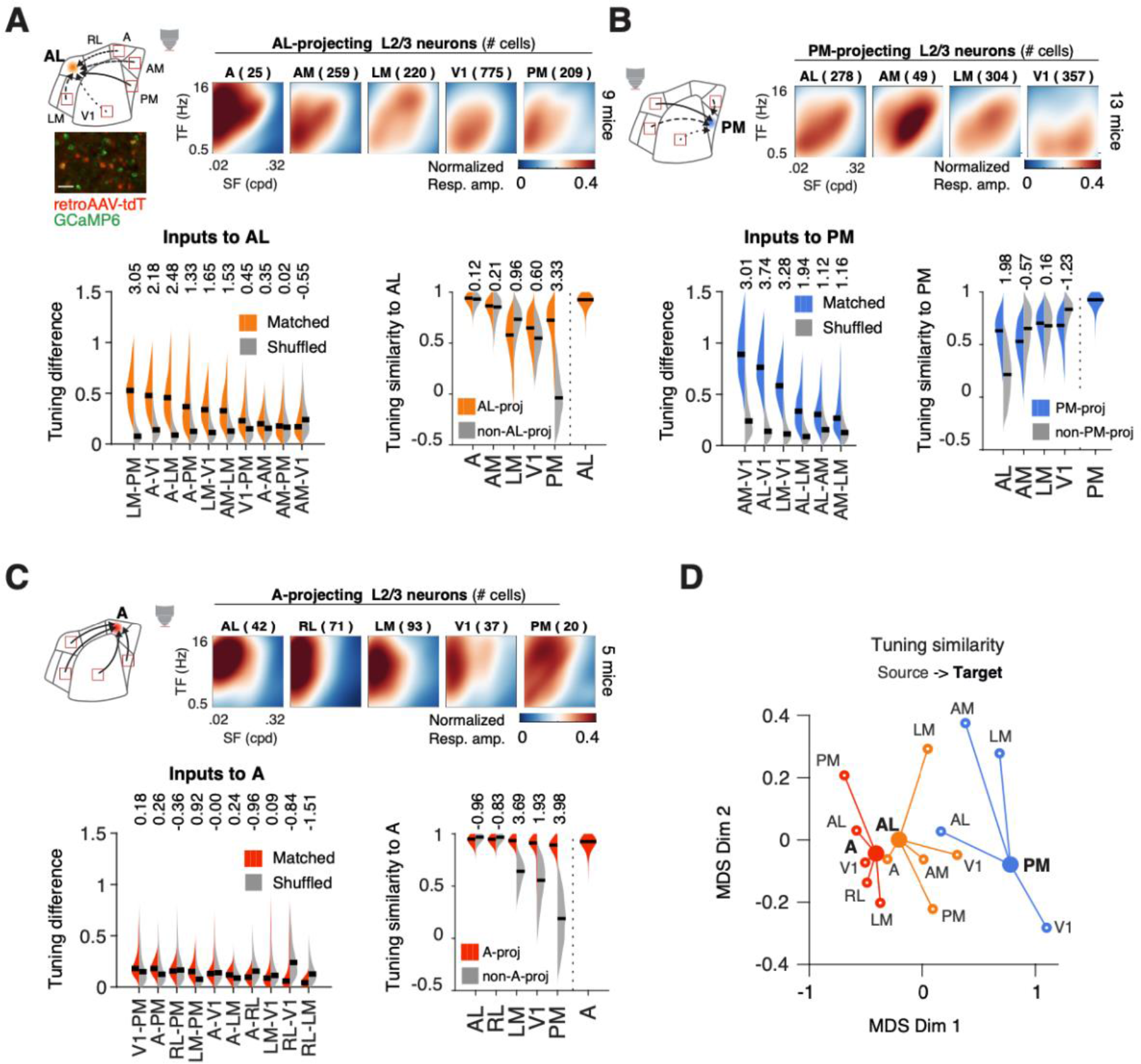
Distinct connectivity rules amongst L2/3 intracortical neurons projecting to HVAs. (A) Diversity and specificity of tuning amongst L2/3 neurons targeting AL. Top left: scheme showing 2-photon imaging of input pathways. Inset at the bottom shows a 2-photon FOV with tdTomato+ retrograde projection neurons and GCaMP6s+ neurons. Top right: summary of average spatiotemporal tuning of retrogradely labeled AL-projecting L2/3 neurons in different areas. Bottom left: Tuning differences between AL-projecting pathways. The differences (colored) were compared to those of the shuffled datasets (gray), and the statistical differences (Cohen’s d) were reported as text at the top. Black horizontal bars: median values of each violin distribution. Bottom right: Tuning similarity of AL with the respective input pathways (AL-proj) and the control group (non-AL-proj). The statistical differences (Cohen’s d) between the two distributions (AL-proj vs. non-AL- proj) are reported as text at the top. As a reference of the within-area variability in AL, the tuning similarity between different subsamples of AL neurons was shown in the rightmost column. Black bars: median values. (B) Same as (A), for input pathways to area PM. (C) Same as (A), for input pathways to area A. (D) 2D-MDS representation of the spatiotemporal tuning similarity between HVAs and L2/3 cortical input pathways. Target HVAs are connected to their respective source areas. MDS: multi-dimensional scaling. See also Figure S3.

Comparing the tuning of projection neurons across areas of origin, we found HVAs receive both diverse and specific visual information from intracortical input pathways. The populations of neurons projecting to AL and PM show diverse tuning, which varies by area of origin (Figure 3A- B, top). In comparison, A projecting neurons are more stereotyped, showing correlated tuning resembling the tuning of A neurons (Figure 3C, top). To quantify diversity amongst input pathways, we computed the distributions of pairwise tuning differences amongst populations of projection neurons, comparing the observed difference with that obtained for shuffled data (Figure 3A-C, bottom left) (see Star Methods). In a subset of projections, the tuning differences amongst neurons projecting to AL or PM deviate from that expected from shuffled data (Figure 3A-B, bottom left; shuffled identities of the projection pathways), indicative of heterogenous visual inputs from distinct visual areas. In comparison, neurons projecting to A did not show such a difference, indicative of more homogeneous visual inputs (Figure 3C, bottom left).

Despite their heterogeneity, the tuning properties of the pathways show an overall bias towards the tuning of their targets. This bias is visible in the multi-dimensional scaling of the pairwise correlation matrix of average tuning (Figure 3D, S3C). To determine whether the similarity of tuning to the target area is due to specific connectivity or to the overall tuning bias in the source area, we compared, using pairwise correlation analysis, the tuning of intracortical projection neurons to the tuning of neurons in the target area (Figure 3A-C, bottom right; S3B, input pathways vs. target areas, Cohen’s d). We found that, for the average tuning, only a subset of pathways shows target-specific connectivity, where the projection neurons differ from the non-labeled neurons showing higher tuning similarity to the target areas (e.g., Figure 3A bottom right, AL-proj vs. non-AL-proj, Cohen’s d). Amongst the input pathways to AL and PM, the reciprocal pathways between AL and PM are the most target-specific (Figure 3A,B bottom right, AL-PM). The other input pathways to AL and PM show little indication of specificity using this metric. Intracortical pathways to A, in contrast, show more target specificity, with projections from V1, LM and PM showing significant average tuning specificity (Figure 3C, bottom right).

To study the population input patterns underlying specific connectivity, we examined the 2- dimensional distributions of peak spatial and temporal frequencies amongst retrogradely (tdT+) and non-labeled (tdT–) neurons in presynaptic visual areas. Similar distributions of peak spatiotemporal frequencies indicate random connectivity, whereas distinct distributions are evidence of specific connectivity. To compare distributions of spatiotemporal frequencies, we used a two-sample two-dimensional Kolmogorov-Smirnov test (2d-KS test).

The analysis of peak spatiotemporal frequencies yielded similar results whereby V1 and LM pathways to AL and A and reciprocal pathways between AL and PM show strong specificity. In contrast, other pathways show comparatively less specificity (Figure 4, S4). Thus, intracortical pathways provide diverse information to HVAs and vary widely in their degree of specificity. Some of the diversity of inputs to HVAs is imputable to target specific projections, while others reflect a functional bias in the source area. Interestingly, bidirectional AL – PM pathways appear to show the highest degree of specificity amongst all probed intracortical pathways.

**Figure 4.**
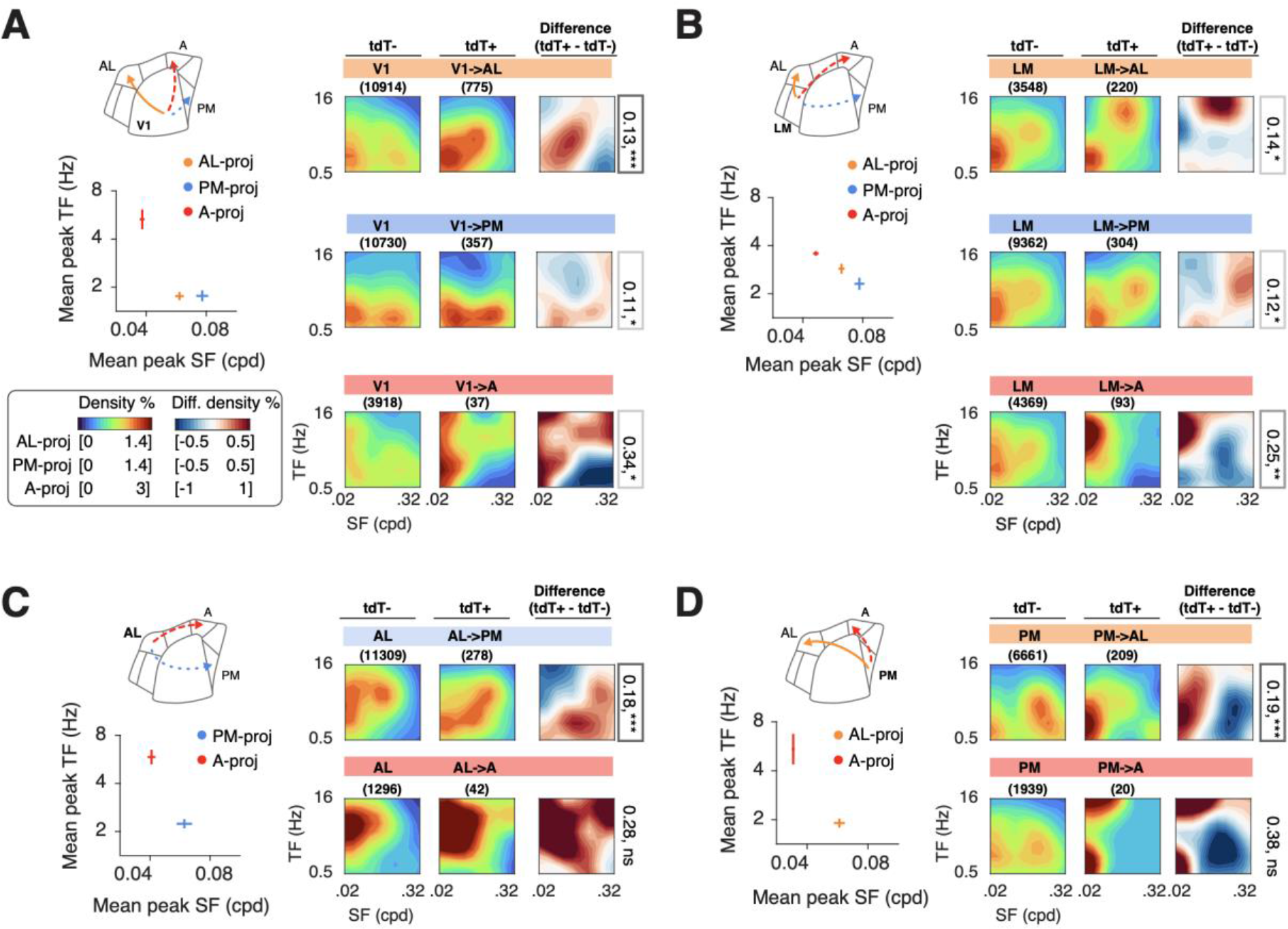
V1 and HVA neurons convey biased, target specific visual information to HVAs. (A) Comparison of the spatiotemporal tuning of V1 neurons projecting to either area AL, PM, or A. Left: Distinct mean peak SF and TF of V1 subpopulations projecting to area AL, PM and A. Means and STD were estimated from the means of bootstrap subsamples of neurons within each group. Right: Density plots showing the distribution of peak spatiotemporal frequencies of V1 L2/3 neurons without (tdT-) and with retrograde labeling (tdT+), and the differences between the two populations. Each row shows the datasets obtained from mice with AL, PM, and A injections, respectively. 2D-KS distances of the distributions of peak spatiotemporal frequencies between tdT- and tdT+ were shown on the right. The significance levels were shown as * and the boxes. p<0.001: ***; p<0.01:**;p<0.05:*; n.s: no significance. Left bottom: color space represents different ranges for different datasets to avoid saturation. Same color spaces for panels A-D. (B) Same as (A), for projection neurons in area LM projecting to either area AL, PM, or A. (C) Same as (A), for projection neurons in area AL projecting to either area A or PM. (D) Same as (A), for projection neurons in area PM projecting to either area A or AL. See also Figure S4.

### L5 HVA neurons show weak target specificity and broadcast similarly tuned visual signals

While HVA neurons in L2/3 can send specific visual information to other HVAs, it is unclear whether the specificity of HVA pathways extends to neurons in other cortical layers. L5 neurons target both cortical areas and deep brain targets hence could abide by distinct connectivity rules (Kim et al., 2015; Lur et al., 2016); however, L5 neurons in V1 were shown to display target specificity (Glickfeld et al., 2013). To examine the specificity of L5 HVA neurons, we used multiplane 2P imaging in retroAAV-CAG-tdTomato injected GCaMP6 mice to image the spatiotemporal visual tuning of AL and PM neurons in L2/3 and L5 (Figure 5A-D, ∼200-500 um deep, 6 mice for ALèPM pathway, 5 mice for PMèAL pathway). The 2P imaging yielded robust neuronal tuning in L2/3 and L5 neurons (Figure S5A). We, therefore, compared the tuning of tdT+ and tdT L2/3 and L5 neurons in the source areas to the tuning of neurons in the target areas.

**Figure 5.**
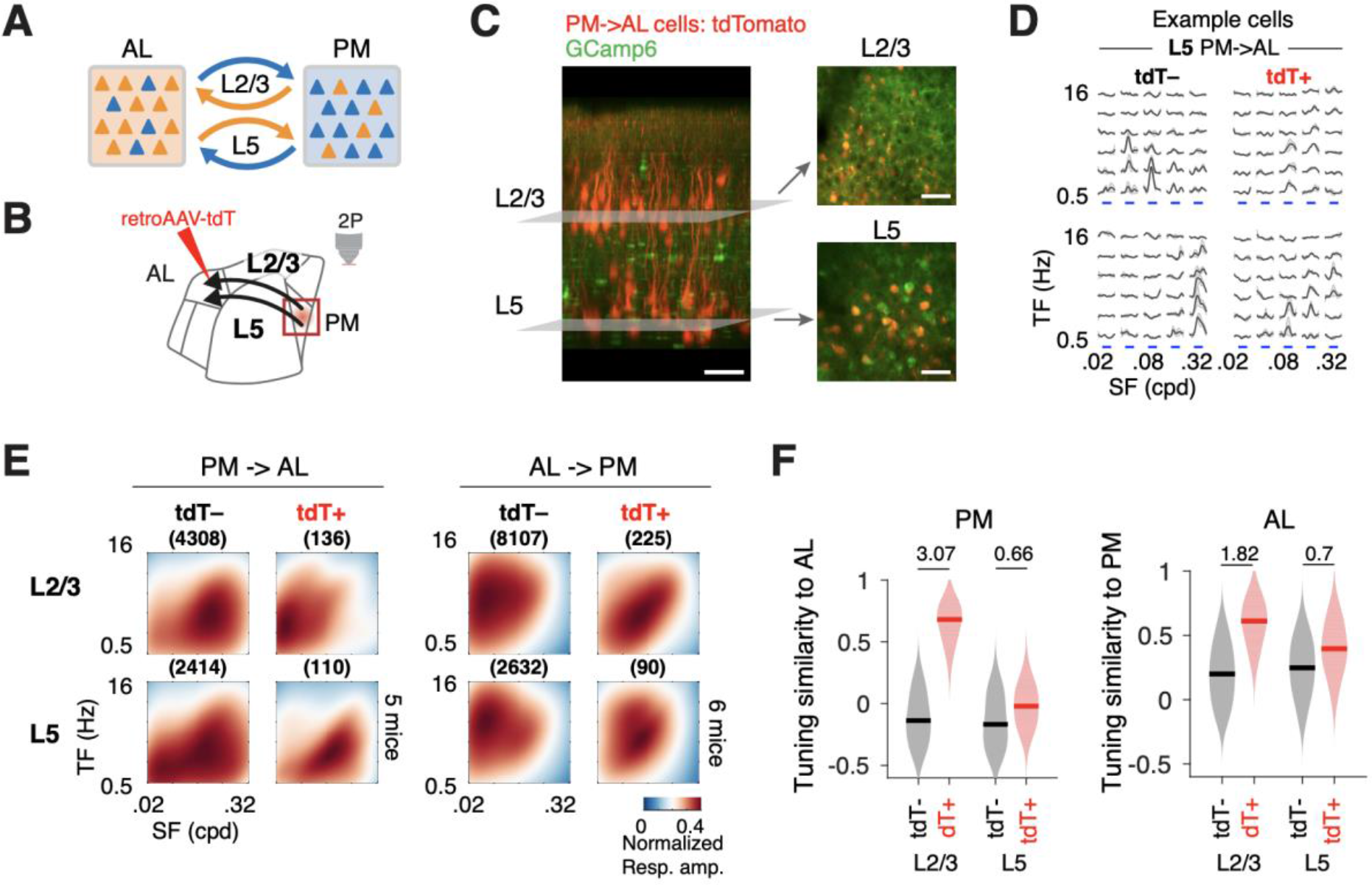
Distinct biases amongst L2/3 and L5 intracortical neurons projecting to HVAs. (A) Scheme showing L2/3 and L5 intracortical neurons mediate communication pathways between AL and PM. (B) Experimental scheme showing retrograde labeling and simultaneous 2-photon imaging of L2/3 and L5 projection neurons. (C) 2-photon volumetric imaging of GCaMP6-expressing neurons in L2/3 and L5 of area PM with retrograde labeling of PM->AL projecting cells. Left image: side-view of the cortex obtained using in vivo volume imaging. Right image: top-view FOVs at L2/3 and L5. (D) Example responses of PM L5 neurons with (tdT+) and without retrograde labeling (tdT-) to spatiotemporal visual stimuli. (E) Average spatiotemporal tuning of tdT- neurons in L2/3 and L5 of PM and AL, and tdT+ neurons mediating reciprocal communication between PM and AL. The numbers of neurons are shown in parentheses. (F) Bootstrap distribution of tuning similarity between the tuning of the subpopulation of neurons (PM or AL) of distinct projection targets (tdT+ vs. tdT-) and laminar positions (L2/3 vs. L5) and the tuning of AL or PM neurons. Blank bars: median values. Pairwise differences are shown as Cohen’s d values at the top. See also Figure S5.

The experiments revealed a lack of specificity in L5 HVA neurons relative to L2/3 HVAs neurons. HVA L2/3 and L5 neurons show similar tuning biases (Figure 5E, tdT–, L2/3 vs. L5; Figure S5B). As seen in the activity of L2/3 neurons, L5 neurons in AL prefer low spatial and high temporal frequencies. In contrast, L5 PM neurons have the opposite bias, preferring high spatial and low temporal frequencies mirroring the tuning preference observed in the activity L2/3 PM neurons (Figure 5E; S5B-C). However, unlike L2/3 HVA-to-HVA projection neurons, which show tuning preferences shifted towards the preferences of their targets (Figure 5E-F, L2/3 tdT+), L5 HVA-to- HVA projection neurons show tuning preferences resembling those of the general population (Figure 5E-F, L5 tdT+). Specifically, while L2/3 tdT+ PM neurons projecting to AL show a shift in spatial and temporal preferences toward the overall preference of AL neurons, L5 PM tdT+ neurons do not. Similarly, the tuning of L2/3 AL (tdT+) neurons projecting to PM show a shift in the direction of the overall tuning preference of PM neurons. This shift is not observed for L5 AL (tdT+) projecting neurons. Thus, while AL and PM L5 neurons show functional specialization, L5 AL and PM pathways do not show the target specificity observed in L2/3.

To determine whether weak specificity is restricted to AL and PM pathways or a general characteristic of HVA L5 pathways, we looked at the visual information conveyed by the axonal projections of L5 neurons in area AL or PM across multiple visual cortical areas (Figure 6A). To selectively label L5 neurons, we injected in transgenic mice with Cre-expressing L5 neurons (Rbp4-Cre mouse line, 4 mice for AL injection, 4 mice for PM injection) AAV that induces Cre-dependent expression of GCaMP8 (AAV-CAG-Flex-GcaMP8m) (Zhang et al., 2023)(Figure 6B). We then used 2P imaging to measure the spatiotemporal tuning of the transfected L5 neurons in AL or PM, as well as the tuning of their axonal projections in the superficial layers (∼20-150 um deep) of distant cortical areas (Figure 6B-C). In this Cre driver line, L5 V1 neurons were shown to display target specificity (Glickfeld et al., 2013). While functionally tuned axons from AL L5 neurons were observed in V1 and all dorsal HVAs, axons from PM L5 neurons to areas RL and A were too sparse to be examined. Putative axonal boutons, the ‘hot spots’ on the axons with high calcium fluorescence fluctuations, were identified (Figure 6C). Stimulus-evoked responses of axonal boutons were used to estimate the spatiotemporal tuning parameters, as for the cellular imaging datasets (Figure 6D).

**Figure 6.**
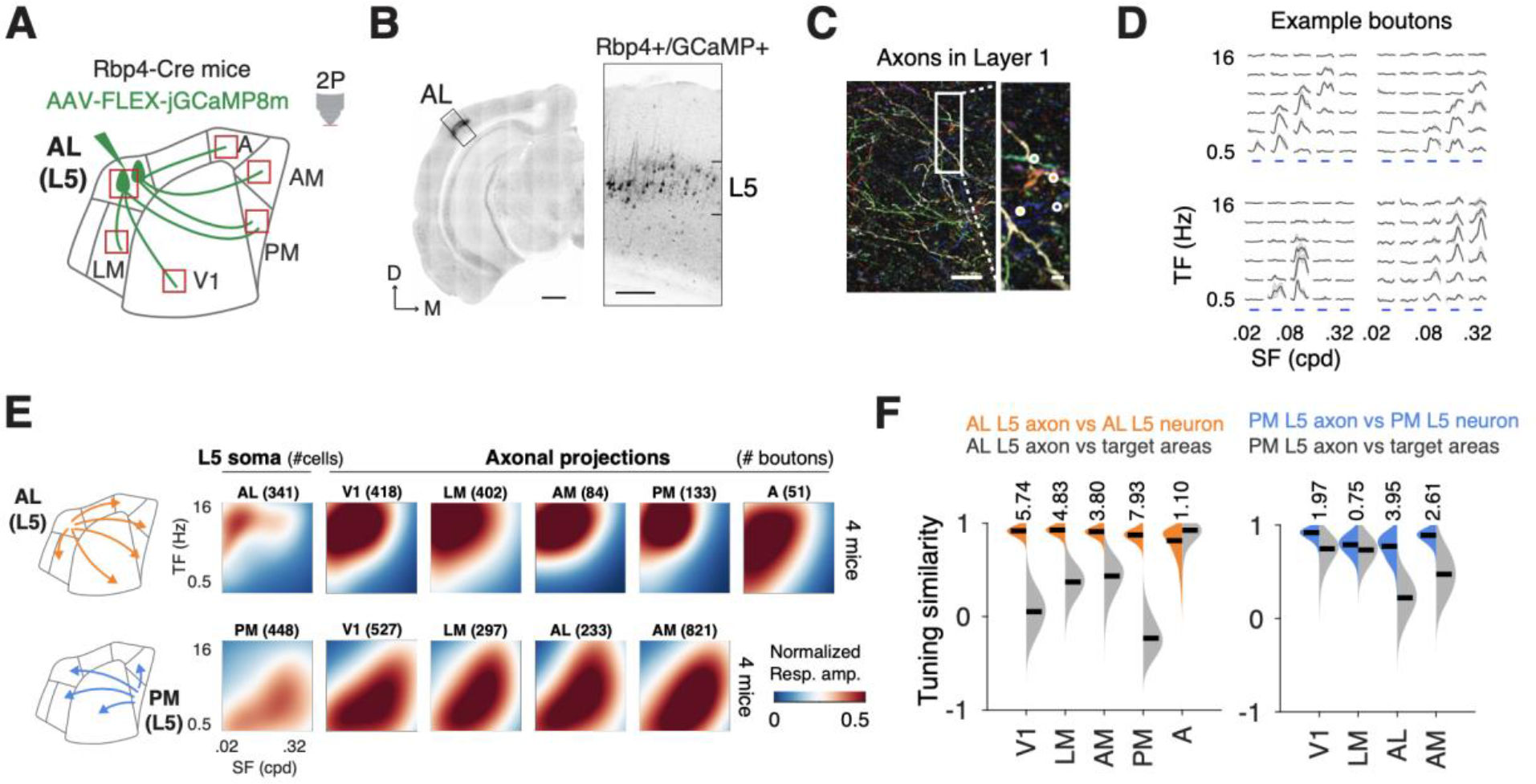
L5 HVA neurons broadcast similar visual information across visual areas (A) Schematic showing the labeling and imaging of axonal projections of L5 neurons. Cre- dependent AAV was delivered to selected HVAs of Rbp4-Cre mice to induce GCaMP8 expression in L5 neurons. Labeled L5 and axonal projections were imaged across cortical areas using 2P imaging (red boxes). (B) Coronal slice showing the laminar location and spread of labeled neurons. Scale bar: 1mm. Right: zoom-in at the injection site (black box). Scale bar: 250um. (C) Example FOV showing axonal projections imaged in cortical layer 1. Axons are color- coded by different spatiotemporal tuning preferences and belong to distinct neurons. Scale bar: 20um. A zoom-in of the white box is shown on the right. White circles show examples of axonal boutons. Scale bar: 2um. (D) Example responses to spatiotemporal visual stimuli of the axonal boutons shown in (C). (E) Average spatiotemporal tuning maps of L5 neurons in AL or PM and their axonal projections in different cortical areas. (A) Bootstrap distribution of tuning similarity of the L5 axons to the L5 source neurons (colored) and the target areas (gray). Black bars: median values. Pairwise differences are shown as Cohen’s d values at the top. See also Figure S6.

Examining the tuning of axonal projections from these areas, we observed that the labeled L5 neurons broadcast highly stereotyped visual information to other visual cortical areas (Figure 6E). Like the cellular data measured in L5 of TRE-GCaMP6s x CaMKII-tTA mice (Figure 5), GCaMP8 labeled L5 neurons in AL and PM show opposite spatiotemporal tuning preferences (Figure 6E, left). However, in all areas, the tuning of axons matches that of neurons in the source areas (Figure 6F, L5 axon vs. L5 neurons), which is not necessarily the case for the target areas (Figure 6F, L5 axon vs. target). Therefore, unlike L2/3 pathways which are functionally target-specific, L5 pathways broadcast similar visual information to recipient areas, potentially contributing to functional diversity.

This observation in the spatiotemporal tuning properties of L5 neuron subsets is quite distinct from that observed when imaging axonal projections labeled by non-layer-specific viral injection (AAV2/1.Syn.GCaMP6f.WPRE.SV40 in AL or PM of C57BL mice) (Figure S6). Furthermore, when looking at the functional specificity of overall axonal outputs from neurons across layers (including both L2/3 and L5) in area AL or PM, we found that both AL and PM (especially AL) outputs are more diverse compared to L5 outputs alone (Figure S6D-G).

### LP neurons carry specific and diverse visual information to HVAs

Although specific, the tuning properties of intracortical pathways cannot easily explain the functional biases of HVAs (e.g., tuning for low SF and high TF in AL, Figure 1G). Overall, the biases in spatial, temporal, and speed preferences are relatively minor compared to the preferences of HVAs (Figure S7A). Therefore, we hypothesized that LP inputs, with their high specific anatomical connectivity (Beltramo and Scanziani, 2019; Bennett et al., 2019; Juavinett et al., 2020), impart specific visual information to HVAs and drive area-specific functional biases.

To investigate this hypothesis, we characterized the visual tuning of LP axonal projections innervating V1 and HVAs (Figure 7A). Using stereotaxic injections of AAV2.1-hSyn-GCaMP6f viral vector in naïve C57/BL mice (5 mice), we induced GCaMP6 expression in LP neurons and their axonal arborizations (Figure 7B; Figure S7B). Using 2P imaging in V1 and HVAs, we recorded LP axons’ responses to visual stimuli of different spatial and temporal frequencies and characterized the tuning of responses. Labeled LP axons were found in V1 and subsets of HVAs. We focused on axonal boutons in the superficial cortex (∼20-150 μm from the pia).

**Figure 7.**
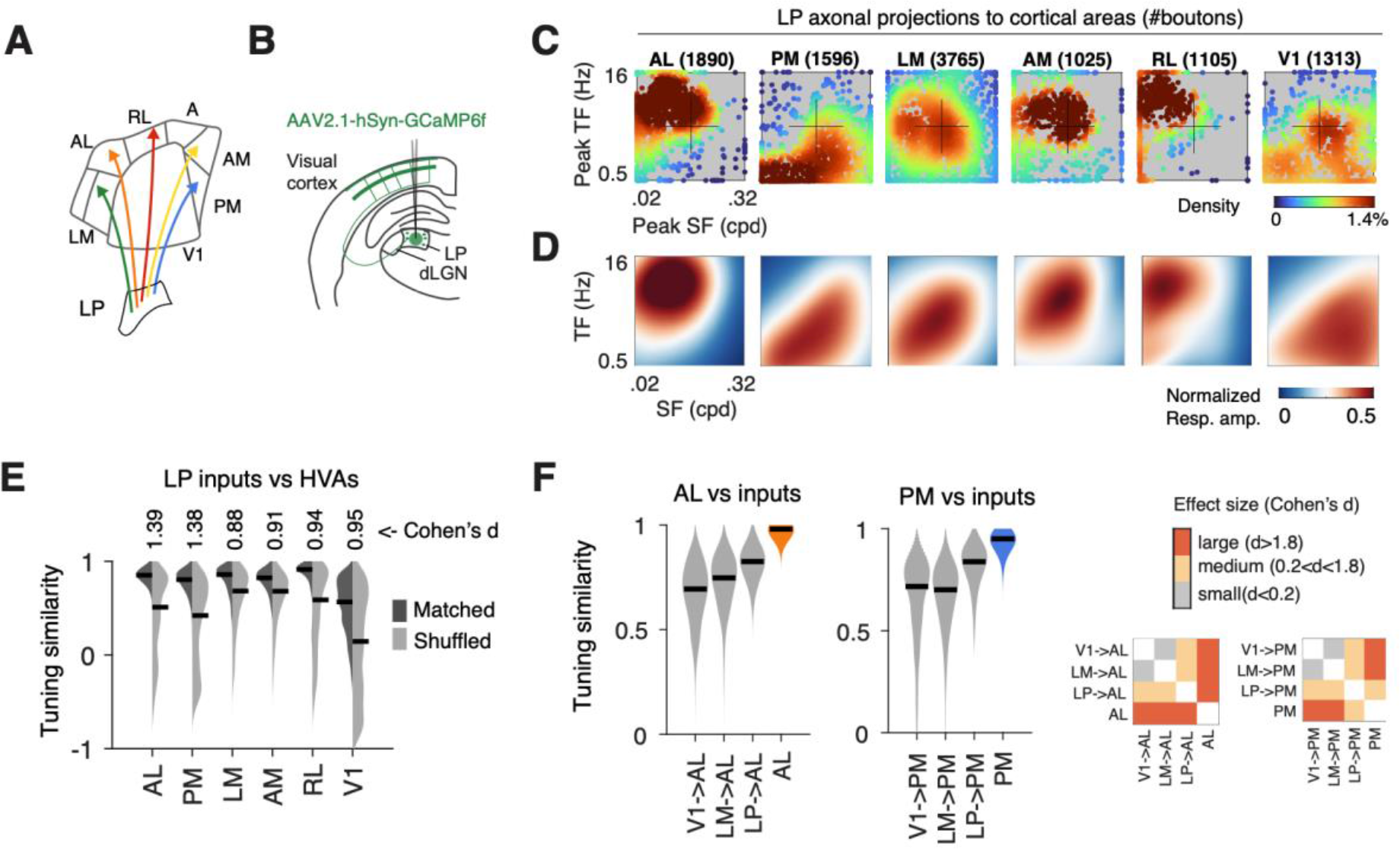
LP neurons convey highly specific, highly diverse visual information to HVAs. (A) Schematic of parallel thalamocortical pathways from the lateral posterior thalamus (LP) to different HVAs. (B) Schematic of labeling and imaging of thalamocortical pathways. LP neurons were labeled with GCaMP6 via the injection of AAV in LP. The axonal projections of LP neurons extending to the cortex were imaged in the superficial layer of different cortical areas using 2P calcium imaging. (C) Distributions of peak SF and TF of axonal boutons of LP neurons in different cortical areas. (D) Average spatiotemporal tuning maps of axonal boutons of LP neurons in different cortical areas. (E) Bootstrap distribution of tuning similarity of LP input pathways to matched cortical targets versus the similarity to shuffled targets. Pairwise statistical differences between matched and shuffled groups (Cohen’s d) are shown at the top. Black bars: median values. (F) Bootstrap distribution of the tuning similarity of different input pathways (from LP, V1, or LM, respectively) to area AL (left) or PM (right). Black bars: median values. Right panes: Matrices showing the pairwise statistical differences between cell groups measured as effect sizes (Cohen’s d). See also Figure S7.

The axonal projections from LP to different HVAs show distinct spatiotemporal tuning (Figure 7C, peak SFTF; 7D, population tuning maps). Whereas the LPèAL pathway shows responses tuned to low spatial and high temporal frequencies, the LPèPM pathway is selective for high spatial and low temporal frequencies. These two pathways encode almost non-overlapping ranges of preferred frequencies. Such functional segregation is observed across higher order thalamocortical pathways, where the LP projections targeting different higher visual areas exhibit widely different spatiotemporal selectivity (Figure 7C-D, S7C).

Furthermore, the tuning of LP axons in HVAs matches the tuning preferences of the target HVAs (Figure 7E, matched vs. shuffled, Cohen’s d), showing pronounced target-specificity. In comparison, LP axons in V1 do not show such a match (Figure 7E, V1), with LP to V1 projections showing functional tuning biases distinct from those of V1 neurons (Figure 7C,D, LP->V1; compare to V1 in Figure 1E). Since projections from LP to HVAs and V1 are considered feedforward and feedback pathways (Miller-Hansen and Sherman, 2022), LP pathways might contribute differently to the visual representation and visual processing in HVAs and V1 with inputs of distinct specificity.

Finally, while L2/3 intracortical and LP pathways carry target-specific information to HVAs, they differ in specificity and might differently shape the visual representations in HVAs. The input pathways showing a higher degree of specificity might drive the biases of visual representations, while less specific but diverse inputs may contribute to the diversity of visual representation. Comparing the target specificity across the main input pathways to HVAs, we found LP pathways are more functionally aligned to HVA populations than intracortical pathways (Figure 7F), suggesting a primary role of higher order thalamocortical pathways in the formation of specialized visual representations in HVAs.

## DISCUSSION

In this study, we characterized the functional organization of lower and higher order intracortical and LP pathways to HVAs. We found HVAs receive diverse inputs from V1, HVA, and LP neurons. However, we uncovered regularities and connectivity rules. First, the density of intracortical inputs to HVAs correlates with the similarity of tuning between visual areas (Figure 2). Second, HVA input tuning correlates with HVA’s preferences and functional heterogeneity (Figure 3-4). Third, HVA output pathways differ widely in tuning homogeneity and target specificity (Figure 5-6). Fourth, LP sends highly differentiated inputs to HVAs that match the differentiated tuning preferences of HVAs.

The results indicate that higher order intracortical and thalamocortical pathways show functionally specific connectivity and may contribute to specialized HVA representations. The results also indicate distinct pathways follow distinct connectivity rules. While intracortical V1, L2/3 HVA, and LP pathways provide specific, biased inputs to HVAs, specific HVA L5 populations broadcast specialized visual information across the visual cortex. This laminar difference in the function projection pathways may reflect input and output pathway specializations. Finally, the results also suggest distinct pathways contribute distinctly to specialized representations. The tuning properties of HVA pathways indicate both intracortical L2/3 and LP pathways may contribute to the tuning specificity of HVAs, with LP playing a prominent role in HVA differentiation. In contrast, L5 pathways that lack target specificity may contribute to the heterogeneity of tuning in HVAs. Further studies are needed to determine the functional implications of these connectivity rules.

### Organization of intracortical visual pathways to and from HVAs

Our data indicate that feedforward intracortical pathways V1 and LM to HVAs are biased and functionally target specific but may only play a limited role in HVA differentiation. L2/3 V1 and LM neurons send biased visual information to AL or PM, consistent with previous studies (Glickfeld et al., 2013; Kim et al., 2018). However, the specificity of V1 and LM to PM pathways is relatively minor (Figure 4A-B), and the strength of the biases is too weak to fully explain AL and PM’s functional biases (Figure 7F, S7A). For area A, although the input from V1 and LM are highly specific (Figure 3), these inputs may be too sparse to be the source of differentiating inputs (Figure 2).

Our data indicate that higher order intracortical visual pathways from HVAs also make specific connectivity. For example, AL and PM L2/3 neurons make specific connections and send biased visual inputs to other HVAs, including to each other, despite their distinct tuning biases (Figure 5, S6). Furthermore, area AL receives substantial inputs from RL, AM, and A (Figure 2C-D) and these inputs have a high specificity in terms of tuning similarity to the tuning of AL neurons (Figure 3A). Area PM also receives inputs from AL and AM. Interestingly, these inputs do not match PM’s biases and thus may contribute to PM’s diversity of tuning. Area A receives precise visual inputs from L2/3 neurons in lower and higher visual areas (Figure 3C). In this case, tuning specificity held despite source areas’ vastly distinct tuning properties (e.g., PM). These properties indicate that in addition to V1 and LM pathways, higher order pathways from HVAs may also contribute to the tuning specificity of HVAs.

In contrast to L2/3 HVA neurons, L5 HVA neurons show a widely distinct connectivity pattern, broadcasting the same specialized visual information across visual cortical areas (Figure 5-6). The more homogeneous tuning of these neurons suggests a distinct functional role, such as increasing functional tuning heterogeneity in postsynaptic populations. Rbp4+ L5 neurons comprise three transcriptional subtypes with distinct laminal positions and projection targets (Harris et al., 2019; Kim et al., 2015). The L5a intratelencephalic (IT) subtype mainly projects within the cortex and could be the primary source of the axons we imaged in distal HVAs (Figure 6). In V1, Rbp4+ L5 neurons send biased visual information to HVAs (Glickfeld et al., 2013). Like L2/3 neurons, L5 Rbp4+ IT HVA neurons receive inputs from a broad yet specific set of visual cortex areas (Yao et al., 2022). Thus, the homogeneity of tuning properties observed is likely not the result of more homogeneous, less diverse long-range inputs.

Altogether, the results on intracortical pathways indicate that these pathways contribute to functional and cell type-specific visual representations. Several subtypes of intracortical projection neurons expressing diverse molecular markers have recently been identified (Tasic et al., 2018). Each shows distinct axonal termination patterns in the target cortical areas (Harris et al., 2019). Divisions of these populations are highly likely since the reciprocal connections between two areas mediated by neurons expressing the same molecular marker can target different layers (Harris et al., 2019). These subtypes might follow distinct wiring rules and send distinct information. Ultimately, multi-dimensional surveys, combining in vivo physiology and anatomy with single-cell transcriptomic profiling of long-range projection neurons, will lead to a better understanding of the genetic and functional logic of long-range connectivity in the cortex.

### Organization of thalamocortical LP pathways to HVAs

Higher order thalamocortical pathways from LP to HVAs show distinct visual tuning and remarkable functional target-specificity, suggesting parallel segregated visual streams that may impart different functional biases to different HVAs (Figure 7). A recent study found LP to AL projections more functionally similar to AL neurons than V1 to AL projections (Blot et al., 2021). Our results are consistent with this finding. That same study, however, reported weak specificity for LP to PM projections. We found a close correspondence between the tuning of LP axons and their HVA targets for five HVAs. Interestingly, the functional correspondence of LP pathways to HVAs was the highest of all pathways we have probed. This result is consistent with a model in which pulvinar pathways that make topographic projections with HVAs (Bennett et al., 2019; Juavinett et al., 2020) impart these areas with their distinct functional and anatomical identity (Murakami and Ohki, 2023; Tohmi et al., 2014).

What might be the origins of visually tuned signals we observed in LP projections to HVAs? LP receives visual information mainly from the superior colliculus (SC) and the visual cortex. From SC, widefield neurons, which are selective for high-spatial and low-temporal frequencies (Gale and Murphy, 2014), preferentially target the posterior LP that connects to ventral HVAs (Beltramo and Scanziani, 2019; Bennett et al., 2019). Hence SC is unlikely to be the source of high temporal frequency information prevalent in anterior dorsal HVAs (Figure 1E). From the visual cortex, both feedforward (V1èLP) and feedback (HVAèLP) pathways were reported to contribute to LP overall activities (Bennett et al., 2019; Blot et al., 2021). LP subpopulations may inherit specific visual selectivity from V1 corticofugal neurons, which get amplified through the reciprocal LP— HVA connections (Bennett et al., 2019). This possibility remains to be tested. In addition, the prefrontal cortex may exert ‘top-down’ modulation of visual tuning of LP neurons via direct or indirect pathways (Bennett et al., 2019; Hu et al., 2019) and facilitate feature-specific visual processing and behavior (e.g., feature-based attention). Future studies are needed to elucidate LP’s input-output connectivity and function in bottom-up and top-down visual functions.

### Distributed and segregated cortical subnetworks

Our results suggest distributed cortical subnetworks specialized for processing specific visual information. Within each area, neurons with similar projection patterns and visual spatiotemporal tuning properties are more likely to interconnect (Cossell et al., 2015; Kim et al., 2018; Ko et al., 2011; Lee et al., 2016). Diverse intracortical projection cell types with distinct projection targets and/or spatiotemporal visual tuning (Figure 4) (Han et al., 2022, 2018; Kim et al., 2020) hence may form multiple segregated local subnetworks that process and transmit specific spatiotemporal information to different visual areas. Across areas, functionally similar neurons in other areas might form distributed interareal subnetworks via recurrent long-range connections. Such organization is suggested to be present between mouse V1 and HVAs. While V1 neurons send target-specific information to area AL (Glickfeld et al., 2013; Kim et al., 2018), AL, in turn, preferentially targets V1 neurons with AL-like tuning (Huh et al., 2018). Similar reciprocal connections are also found between V1 and PM (Huh et al., 2018) and might exist amongst HVAs. Moreover, local subnetworks in different areas can be interconnected via the collateral axonal projections of intracortical cells (Han et al., 2018). Such recurrent subnetworks might permit signal amplification and prolonged information integration windows (Li et al., 2013a, 2013b; Lien and Scanziani, 2013). Future studies are necessary to determine whether neurons in different cortical areas form monosynaptic reciprocal loops and determine the interaction between functional subnetworks within and between cortical areas.

Our data also suggest different HVAs differentially partake in distributed cortical subnetworks. Areas PM and A encode essentially non-overlapping visual information and participate in separate subnetworks as they avoid connecting (Figure 2) and receive segregated inputs from distinct subsets of areas and different subpopulations in the common source areas (e.g., ALè PM and ALèA; Figure 3,4). Such anatomical segregation constrains the information exchange between the two areas and supports function segregation. Area A might be primarily involved in a multi-area subnetwork in the dorsal cortex for processing fast visual motion. It receives profound inputs from adjacent anterior dorsal areas, showing a strong preference for fast motion (Han et al., 2022). These short, dense connections between anterior areas may provide a neural substrate for transmitting information about fast visual motion and may support the proposed roles of these areas in animals’ exploratory behaviors (La Chioma et al., 2019; Zhuang et al., 2017). In the meantime, area AL may participate in multiple distributed subnetworks with diverse AL subpopulations connecting to distinct neuronal populations or distinct areas.

## Conclusion

This study uncovered cell-type and area-specific logic of intracortical and thalamocortical pathways through a detailed characterization of the anatomical and functional organization of the input and output of higher visual cortical areas. Highly specific pathways (e.g., higher order thalamocortical pathways, L2/3 pathway to area A) can impart functional biases to HVAs. In contrast, less specific inputs (e.g., L2/3 and L5 intracortical pathways to AL and PM) convey diverse information and can contribute to the diversity of visual representations in HVAs. Further studies are needed to study the implications for visual representations and behavior.

## STAR METHODS

### EXPERIMENTAL MODEL AND SUBJECT DETAILS

#### Mice

All experiments were approved by the Animal Ethics Committee of KU Leuven. Three mice genotypes were used: naïve C57BL/6 mice, Rbp4-Cre mice [Tg(Rbp4-cre)KL100Gsat, MGI:4367067] (Harris et al., 2019), and CaMKII-GCaMP6 mice [B6;DBA-Tg(tetO- GCaMP6s)2Niell/J x B6.Cg-Tg(Camk2a-tTA)1Mmay/DboJ; The Jackson Laboratory stock No. 024742 and 007004] (Wekselblatt et al., 2016). The mice were single-housed in an enriched environment (cotton bedding, wooden blocks, and running wheel), with 12h-12h light-dark cycle, 19–21 degrees cage temperature, and 30–70% humidity.

#### Craniotomy

Standard craniotomy surgeries were performed to gain optical access to the visual cortex (Goldey and Andermann, 2014). Mice between 2 and 3 months old were anesthetized with isoflurane (2.5%–3% induction, 1%–1.25% surgery). A custom-made titanium frame was attached to the skull, and a cranial glass window was implanted in the left hemisphere over the visual cortex. In five mice, a 5-mm cranial window was centered over V1 (3.10 mm lateral from lambda, 1.64 mm anterior of the lambdoid suture). Buprenex and Cefazolin were administered postoperatively when the animal recovered from anesthesia after surgery (2 mg/kg and 5 mg/kg, respectively, every 12 hours for 2 days). All mice were habituated at least 3 days before data acquisition.

#### Viral injection

Long-range connectivity was investigated by injecting anterograde or retrograde neural tracers into the cortex or LP. Anesthesia and postoperative treatments were delivered as described in the above section. Visual cortical areas were identified using retinotopic mapping and widefield calcium imaging for GCaMP6 reporter mice for widefield flavoprotein imaging for non-GCaMP6 mice (see details below, Retinotopic mapping). In a typical viral injection, a glass injection capillary (∼15-25 µm internal tip diameter) was first loaded with the viral vector. Then the capillary tip was slowly lowered to the desired coordinates. After 10 min of recovery, viral vectors were slowly pulse-injected (25nl per min, Nanoject II, Drummond Sci.), followed by 10 min recovery before the capillary was slowly retracted from the brain. In order to induce specific expression of GCaMP in L5 neurons, a Cre-dependent adeno-associated virus-based viral vector (AAV1-CAG-FLEX- jGCaMP8m-WPRE, Addgene 162381-AAV1) (Zhang et al., 2023) was injected into selected visual areas (AL or PM, 50nl at 500 um depth) of Rbp4-Cre mice. To transfect excitatory neurons across cortical layers with GCaMP, an AAV-based viral vector (AAV2/1.Syn.GCaMP6f.WPRE.SV40; University of Pennsylvania Vector Core) was injected into selected areas (AL or PM, 50nl each at 500 and 200um depths) in naïve C57BL6 mice. To label intracortical projection neurons projecting to selected cortical areas, a retrograde AAV viral vector (retroAAV-CAG-tdTomato, Addgene: 59462-AAVrg) (Tervo et al., 2016) was injected into selected areas (AL, PM or A, 20nl each at 200 and 500 um depths) of CaMKII-GCaMP6 mice. To label LP neurons with GCaMP, anterograde AAV constructs (AAV1.CAG.GCaMP6f.WPRE.SV40, Addgene: 100836-AAV1) were injected into LP (1.90 mm posterior, 1.70 lateral from bregma, at the depth 2.50 mm) in C57BL6 mice. For inspection of the expression, images were taken using 1-photon and 2-photon imaging (see the following sections). The tdTomato expression increases over time and becomes clearly visible one week after injection. GCaMP6 expression in cell bodies and axons was visible three weeks after injection of AAV vector with a Syn promotor and two weeks after injection for AAV constructs with a CAG promotor.

#### Histology

For histology, mice were perfused with 25 ml 0.1mM PBS and then 25 ml of 4% paraformaldehyde (PFA, Invitrogen). The whole brain was extracted and stored overnight in 4% PFA. A series of coronal or sagittal sections (100um thick) were obtained using a Leica VT100S vibrating blade microtome and stored in PBS with 0.5 mg/mL sodium azide (Sigma). An immunostaining process was performed to enhance the signal of GCaMP6 of axons. Slices were blocked with PBS supplemented with 3% donkey serum (Invitrogen) and 0.3% Triton X-100 (Sigma) for 1h and incubated overnight at 4 °C with the primary antibody (1:1000 anti-chicken, polyclonal, Thermo Fisher Scientific). Slices were rinsed with PBS/0.1% Triton X-100 three times and then incubated 2 h at room temperature with the secondary antibodies (1:800 Alexa 488 donkey anti-chicken, Immuno-Jackson). Sections were again rinsed three times with PBS/0.1% Triton X-100 and then twice in PBS. The sections were next mounted on a Superfrost slide (Thermo Scientific) and dried using a brush before adding histology mounting medium with DAPI (Fluoroshield with DAPI, Merck). Confocal images were acquired using a Zeiss LSM710 confocal microscope controlled by the Zen Black software (Zeiss).

#### Visual stimulation

Visual stimulation was performed as previously reported (Han et al., 2022). A 22-inch LCD monitor (Samsung 2233RZ, 1680 by 1050-pixel resolution, 60 Hz refresh rate, brightness 100%, contrast 70%) with a mean luminance of 54–80 cd/m^2^ (from edges to center) was positioned 18 cm in front of the right eye, covering 100 by 80 degrees of the visual field (0 to 100 deg in azimuth from central to peripheral and -30 to 50 deg in elevation from lower to upper visual field). Visual stimuli were generated in MATLAB (The Mathworks, Natick, MA) and presented using PsychoPy2 and custom Python code. A spherical geometric correction was applied to the stimuli to define eccentricity in spherical coordinates.

To activate cortical neurons with diverse tuning for orientation features and visual motions (Han et al., 2022; Niell and Stryker, 2008), three sets of visual stimuli were used: isotropic noise stimuli (ISO) containing non-oriented textures, anisotropic noise stimuli (ANISO) containing elongated textures, and drifting gratings. The ISO and ANISO stimuli were generated by applying parametrized filters to random pink-noise fields as previously described (Han et al., 2022). Narrow bandpass filters (bandwidth: 1 octave) for spatial and temporal frequencies determine the scale and flickering frequencies of the textures. The orientation bandwidth determines the length/width ratio of the textures. Isotropic noise stimuli, spectrally band-passed noise with no angular filter, resemble clouds of moving dots (Figure S1A, ISO stimuli icons). A narrow orientation bandwidth (15 deg) creates elongated textures (Figure S1A, ANISO stimuli icons). While isotropic textures contain no global motion, oriented textures were designed to rotate slowly (45 deg per second, clockwise) to activate neurons tuned to different orientations. The drifting gratings (Figure S1A, gratings icons) differ from the noise stimuli as they contain coherent visual motion, a key component to activate specific neuronal populations (Sit and Goard, 2020). For the drifting gratings, each spatiotemporal frequency combination was presented as gratings moving in four cardinal directions (up, down, nasal, temporal, 1s for each direction, no blank in between).

To probe the neuron’s selectivity for spatial and temporal frequencies, the three visual stimulus sets, each consisting of 30 combinations of spatial frequencies (0.02, 0.04, 0.08, 0.16, 0.32 cpd) and temporal frequencies (0.5, 1, 2, 4,8,16 Hz), were displayed to the mouse. Each frequency combination was presented for 4 seconds, intertwined with 4-second equiluminant gray screen. All combinations were presented in random orders, for 4 repeats. In each repeat, a unique set of ISO and ANISO noise stimuli was generated using distinct random seeds.

Cortical neurons showed differential preferences for isotropic, anisotropic noise, and moving grating stimuli. In all L2/3 neurons detected across areas, 41% of neurons showed reliable stimulus-evoked responses to at least one stimulus set (75th percentile cross-trial correlation > 0.3). The three stimulus sets activated overlapping yet distinct populations of neurons (Figure S1G). Gratings showed the best capability to drive neuronal responses in terms of the fraction of activated neurons and response amplitudes (Figure S1D,E). The stimuli driving the strongest responses were deemed a neuron’s preferred stimuli. The corresponding responses were used for spatiotemporal tuning analysis (see below, Tuning analysis).

#### Two-photon functional and structural imaging

For 2-photon imaging, a 920-nm femtosecond laser beam (Newport MaiTai DeepSee) was raster-scanned using galvo and resonant scanners (Cambridge 6215H and CRS 8K). With 16x lens (NA = 0.8, Nikon), activity from neurons was recorded typically at 100∼300 µm deep below pia for L2/3 neurons, 350∼500 µm deep for L5 neurons, 200∼500 µm deep for simultaneous imaging of L2/3 and L5 neurons. Axonal projections’ activity was typically recorded at 20∼150µm deep with 2x digital magnification. GCaMP6 fluorescence was collected using a bandpass filter (510/84 nm, Semrock) and GaAsP hybrid photodetectors (H11706-40, Hamamatsu), and images were reconstructed and acquired using Scanbox (version 4.0, https://scanbox.org). Individual imaging planes (720x512 pixel per frame, 1249 by 1067 um field-of-view) were collected at different depths. Simultaneous multiplane imaging was achieved by rapidly changing the focus with an electrically tunable lens (EL-10-30-TC, Optotune AG; staircase mode) from 100 to 300 um depth and varying the laser power (from ∼50 to 120 mW). Images were acquired from 3 or 4 evenly spaced planes at 10.33 and 7.75 Hz, respectively. Blackout material (Thorlabs) blocked stray light from the visual display entering the light collection path.

High-resolution structural image stacks of retrogradely labeled cells in the cortex were obtained by collecting two-photon cellular images every 5 um (controlled by motorized stages, averaged over 50 frames per section) from the pia to 500 um depth in the cortex. In addition, red fluorescence of tdTomato was collected using a bandpass filter (607±35 nm, Semrock).

#### Widefield fluorescent Imaging

Widefield calcium and flavoprotein imaging was conducted using a widefield in vivo microscope (Neurolabware LLC), performed with blue excitation light (479nm LED, 469/35 nm bandpass filter, Thorlabs) through a low magnification (2x) objective lens (NA = 0.055, Mitutoyo) and green emission light (498 nm dichroic, 525/39 nm filter, Semrock) collected with an EMCCD camera (QImaging EM-C2, Teledyne Photometrics; 1,004 by 1,002 pixels with 2 by 2 binning) at a rate of 5 frames per second (fps).

Widefield retrograde labeling imaging was performed with a fluorescent microscope with 1x objective. A green light beam from a halogen lamp passing through an excitation filter was used for illumination. The red fluorescence was collected with a CCD camera (PCO Pixelfly 1.3 SWIR) and acquisition software (PCO Camware). The acquired images were filtered with a bandpass kernel using ImageJ.

#### Retinotopic Mapping

Visual cortical areas were identified using retinotopic mapping as described previously (Han et al., 2022). Briefly, we used a circular visual stimulus (“circling patch”) rotating around the visual display center or a rectangular bar (“traveling bar”) translating along the horizontal or vertical axes of the display. The circling patch stimuli is a small patch of isotropic noise stimuli (0.08 cpd, 2 Hz, 20 deg in diameter) circling along an elliptic trajectory on the display (azimuth: 10 to 90 deg; elevation: -20 to 40 deg; 20 sec per cycle with 20 repetitions). The traveling bar stimuli comprised a narrow band (13 deg wide) of isotropic noise stimuli (0.08cpd, 2Hz) sweeping across the screen in 4 cardinal directions (10 sec per sweep, 20 repetitions). The phase maps were generated by calculating the stimuli’s phase/position, evoking maximal neural responses for each image pixel. Each area has a complete representation of the circling patch’s elliptic trajectory, resulting in a pinwheel-like pattern. The traveling bar stimuli generated azimuth and altitude phase maps, which were used to identify the reversals of retinotopic maps between areas. Areas are identified according to the common delineation (Wang and Burkhalter, 2007; Zhuang et al., 2017).

#### Data Analysis

All subsequent data analysis was performed in MATLAB (The Mathworks, Natick, MA).

#### Anatomical connectivity analysis

To determine the degree to which widefield fluorescence intensity of retrograde labeling (tdTomato) reflects the number of labeled neurons, we measured the density of labeled populations using 2-photon structural imaging (from pia to ∼ 500um deep). We found a high correlation between the two measurements (Figure S2). Regions of high fluorescent intensity contain dense retrograde labeling of projection neurons, distributed in superficial and deep layers, whereas low-intensity regions contain much sparser labeled neurons (Figure S2C). In addition, the average intensities of one-photon images and corresponding two-photon volumes were highly correlated (Figure S2D-E), permitting the use of the total fluorescent intensity of each cortical area as a metric of the total number of retrogradely labeled neurons.

To determine the relative strengths of intracortical input pathways to a given HVA, we calculated, from the widefield fluorescent images, the total fluorescent intensity (sum of all pixels) of retrograde labeling of individual visual cortical areas (*I*_*x*_) that were identified using retinotopic mapping. The anatomical connectivity strength of pathway x is calculated as:

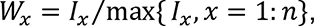

Where *n* is the total number of retrogradely labeled areas.

#### Cellular and axonal calcium imaging data

Imaging processing and analysis were described in our previous study (Han et al., 2022). To correct x-y motion, 2P images for all experiments during the imaging sessions were registered to a common reference image (by registering and averaging 1200 frames from the center of the session). Regions of interest (ROIs) of active neural cell bodies were detected by computing local correlation (3-by-3 pixels neighborhood, threshold at correlation coefficients > 0.9; customized MATLAB software) and identifying spatially connected pixels, selecting ROIs with near-circular shapes (maximum length/width aspect ratio < 2). Active axonal boutons were detected with a similar routine with selection parameters adjusted for small ROIs (area size 4-8 pixels). ROIs fluorescence time courses were generated by averaging all pixels in an ROI, subtracting the averaged neuropil signals in the surrounding ring (morphological dilation, ring size matched to the ROI, non-overlapping with neighboring ROIs) and correcting for slow baseline drift. Raw calcium time courses (ΔF/F_0_) were expressed as fractional changes to the baseline fluorescence.

#### Selection of responsive and reliable cells and axonal boutons

Our previous study described the selection process (Han et al., 2022). A cellular or axonal ROI was classified as responsive if the median time courses computed across trials showed a response peak with magnitude > 3x standard deviation of the pre-stimulus activity (averaged > 2 seconds) over a continuous period > 1 second for at least one stimulus condition. Cells/boutons responding to the visual stimulus offset but not during visual stimulus epochs were excluded from the analysis. ROIs were selected based on a response reliability index, defined as the 75 ^th^ percentile of the cross-trial correlation coefficients of the de-randomized response time courses. ROIs with a reliability index > 0.3 were selected for further analysis. Neurons/boutons’ response amplitudes were calculated as the average ΔF/F_0_ throughout the stimulus epochs.

#### Spatiotemporal tuning analysis

Responses were described by two-dimensional elliptical Gaussian functions (Andermann et al., 2011; Priebe et al., 2003):

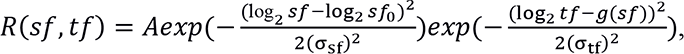

where *A* is the peak response amplitude, *sf0* and *tf0* are the peak spatial and temporal frequencies, *σsf* and *σtf* are the spatial and temporal frequency tuning widths, and

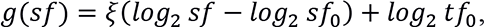

where *ξ* represents the speed-tuning index that captures the slant in the spatiotemporal frequency space. Model parameters were estimated by searching for the model fit with least squared errors (MATLAB *lsqcurvefit* function). Peak speed is the ratio between peak temporal frequency and spatial frequency.

#### Comparison of spatiotemporal peak frequencies distributions

A two-sample two-dimensional Kolmogorov-Smirnov test (2d-KS test) (Peacock, 1983) was used to compare the distributions of neuronal populations in the spatiotemporal frequency space (Figure 1G-H, 4A-D, S4). The yielded distance and significant level (p-value) were reported.

#### Hierarchical bootstrapping

To measure the average functional properties of visual areas of pathways with the consideration of inter-animal variability, we constructed hierarchical datasets using a hierarchical bootstrapping approach (Han et al., 2022). Each hierarchical dataset consisted of 1000 random subsamples (with replacement), each of which consisted of equal numbers of data points from equal numbers of animals (typically, 50 data points per mouse, 5 mice per group). Then the metrics of interest for each subsample were calculated, then the means and distributions of the metrics across subsamples were obtained. All the following analyses used hierarchical bootstrapping.

#### Average tuning similarity

The overall functional bias of a group of neurons was measured as the bootstrap average spatiotemporal tuning map across neurons. Briefly, the average of neurons’ peak-normalized SFTF maps in each subsample was obtained, and the means across subsamples were defined as the bootstrap average SFTF tuning maps (Figure 1E, 3A-C top, 5E, 6E, 7D, S6D). Average tuning similarity between areas or pathways was calculated as pairwise Pearson’s correlation coefficients between bootstrap average SFTF tuning maps of the compared neuron groups (Figure 1F, S3C).

#### Comparison of population tuning similarity

To determine whether the functional similarity between a pair of neuronal populations (i.e., pathways or areas) differs from other pairs, we compared the distributions of functional similarity (obtained using hierarchical bootstrapping) of the two pairs. To do so, we repeatedly selected random subsets of neurons from each cell group, calculated the average SFTF tuning maps, performed pairwise Pearson’s correlation, and compared the distributions of correlation coefficients across bootstraps. For example, to determine whether the similarity between populations A and B is higher than that between populations C and D, hierarchical bootstrap subsamples for each population were generated (subsample A-D), and the averages of neurons’ peak-normalized SFTF maps in each subsample were calculated (map A-D). To obtain the distributions of tuning similarity, referred to as population tuning similarity, between populations A and B, we calculated the Pearson’s correlation coefficients between maps A and B (*r*_AB_) in each subsampling. We examined the distributions of *r*AB across subsamples. Tuning difference is 1 minus the correlation coefficient (Figure 3A-C, bottom left). Then Cohen’s d was used to report the differences between the distributions of *r_AB_* and *r*_CD_ (Figure 3A-C bottom panels, 5E, 6F, 7E- F, S3B, S6E-G). d<0.2 means small difference, 0.2<d<1.8 means medium difference, d>1.8 means large differences.

#### Comparison of peak frequencies

To determine whether neuronal populations projecting to different areas have different functional biases (Figure 3G), the mean and standard error of the mean of each population were obtained using hierarchical bootstrapping. Briefly, the mean peak spatial and temporal frequencies (*μ*_*TF*_, *μ*_*SF*_) of each random subsample were obtained, and the means and standard deviations of *μ*_*TF*_ and *μ*_*SF*_ across subsamples (that is the mean and standard error of the mean of the population) were obtained and used to calculate the differences between populations.

## Acknowledgements

We are grateful to Ben Vermaercke for supporting the imaging setup and image processing, Asli Ayaz, and Karl Farrow for feedback on earlier versions of the manuscript. This work was supported by Neuro-Electronics Research Flanders (V.B.), FWO grants G0D0516N (V.B.), G0C1220N (V.B.), and KU Leuven Research Council grant C14/16/048 (V.B.).

## Author contributions

Conceptualization: X.H. and V.B.; Methodology: X.H. and V.B.; Software: X.H.; Investigation: X.H.; Data curation: X.H.; Formal Analysis: X.H. and V.B.; Visualization: X.H. and V.B.; Writing—original draft: X.H.; Writing—review & editing: X.H. and V.B.; Funding acquisition: V.B.; Resources: X.H. and V.B.; Supervision: V.B.

## Supplementary figures

**Figure S1.**
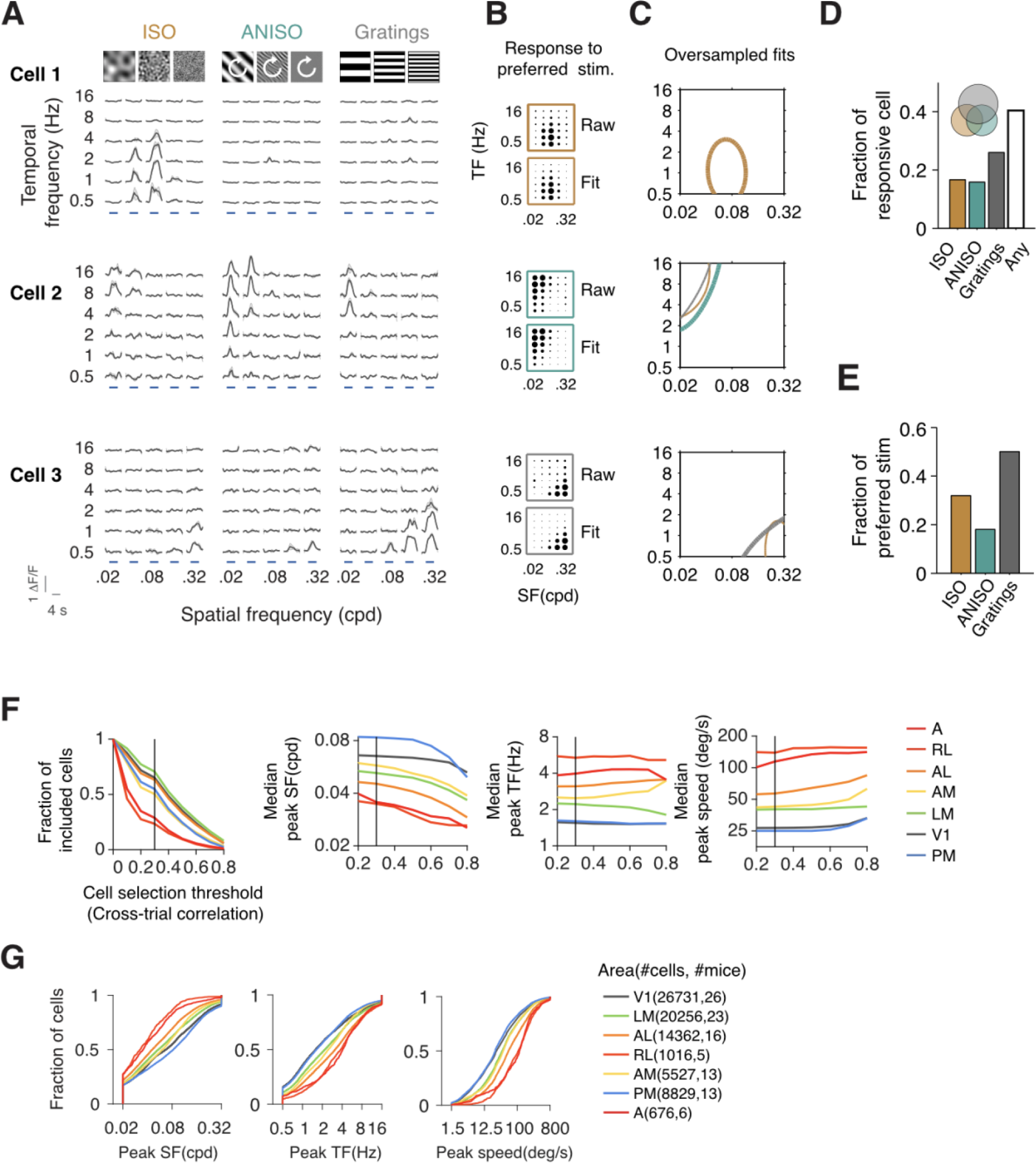
Characterization of spatiotemporal tuning of HVA neurons using noise and gratings stimuli, related to Figure 1. (A) Visual responses (traces of calcium fluorescent intensity) of three example cells (rows) to three spatiotemporal visual stimuli (from left to right: isotropic noise, anisotropic noise, drifting gratings). The same color code for the three stimuli is used throughout panels A-E. (B) Dot plots showing mean response amplitudes (raw) and the model fits of the strongest responses to the three stimuli. (C) Contour plots showing the fits (at 50% peak amplitudes) of responses to noise and gratings visual stimuli. The fits with the strongest response amplitudes are shown with thicker contours. (D) Bar plot showing the fraction of neurons with reliable visual responses to each stimulus set and to any of them. (E) Bar plot showing the fraction of neurons that prefer isotropic noise, anisotropic noise and drifting gratings, respectively. (F) Impact of the cell selection threshold (minimal cross-trial variability) on the fraction of included neurons, median peak spatial frequency, median peak temporal frequency, and median peak speed (from left to right). The threshold was set to 0.3 in the current study. (G) Cumulative distributions of peak spatial frequency, peak temporal frequency and peak speed of L2/3 neurons in different visual areas.

**Figure S2.**
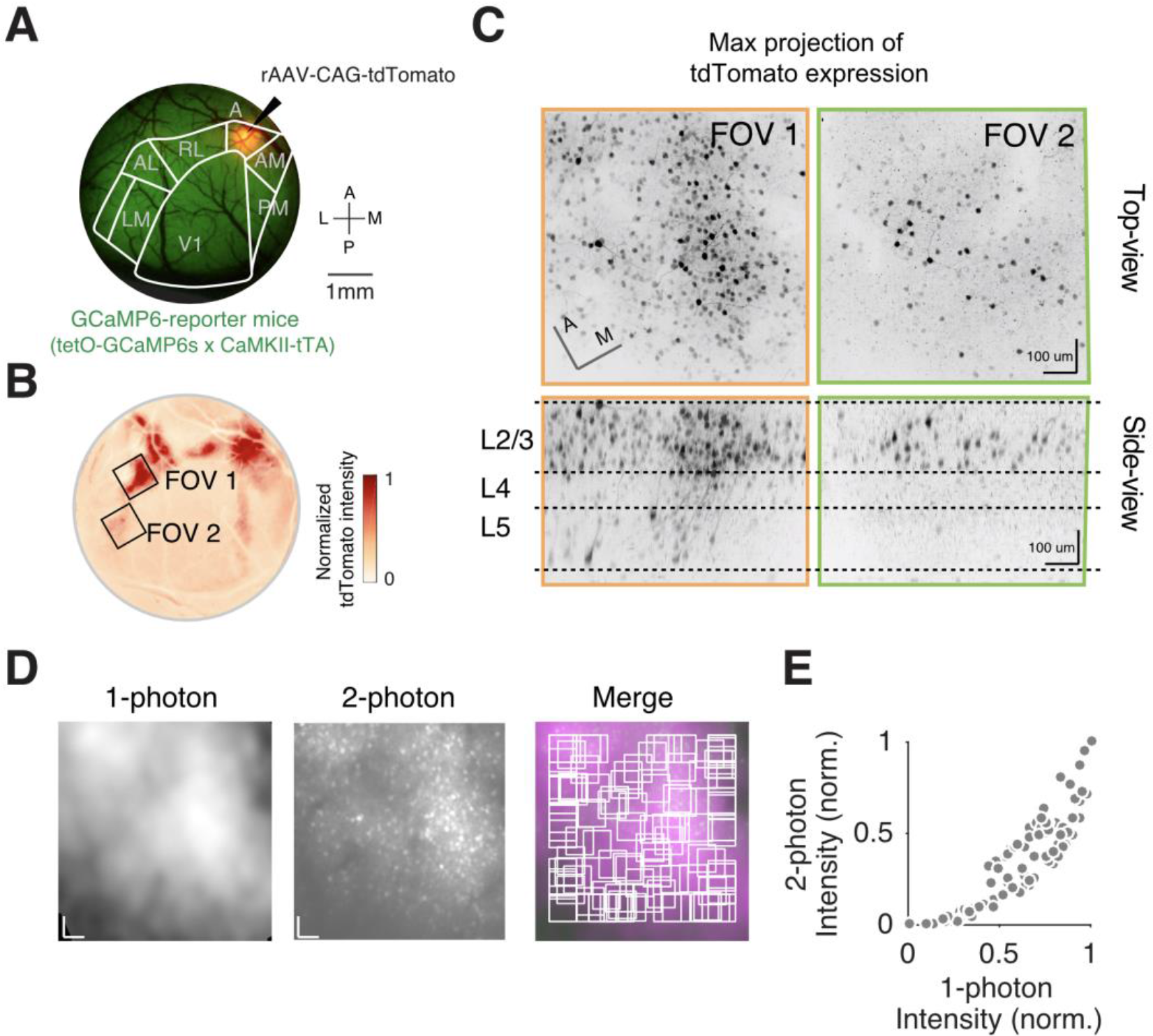
Mesoscopic fluorescent intensity of retrograde labeling broadly correlates with the density of retrogradely labeled neurons, related to Figure 2. (A) Example widefield fluorescent image of a cranial window of a GCaMP6-mice with the injection of retrograde AAV in area A. White contour: area delineation. (B) Cortical input maps as a filtered image of the raw image shown in (A). Two FOVs (black boxes) with distinct intensity levels were further inspected with 2P volumetric imaging (results shown in panel C). (C) Max projections of the tdTomato expression showing the top-view and side-view of FOV1 and 2. (D) Comparing signals of 1P FOV and corresponding 2P FOV. A 2D Gaussian filter was applied to the 2P FOV to approximate the effect of light scattering on 1P imaging. The average intensities of different regions of the FOV are plotted in (E). (E) Scatter plot showing the correspondence of the intensity of retrograde labeling measured using 1P or 2P imaging.

**Figure S3.**
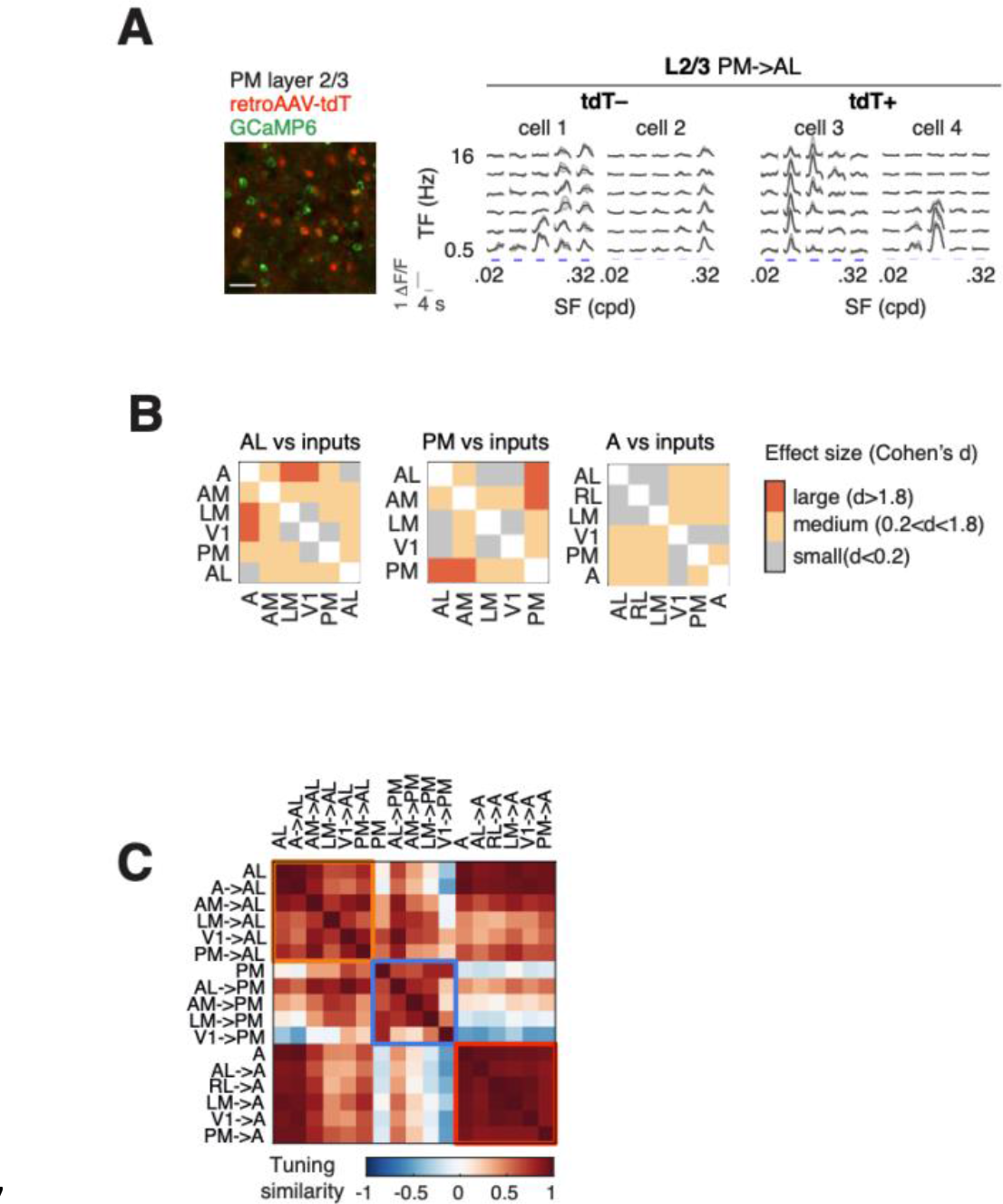
L2/3 intracortical pathways are functionally target specific, related to Figure 3. (A) Left: example FOV showing GCaMP6-expressing L2/3 neurons (green) in area PM with a retrogradely labeled subpopulation that projects to area AL (red and yellow). Scale bar: 100um. Right: example responses to spatiotemporal visual stimuli. (B) Matrices showing Cohen’s d between the target specificity of input pathways to areas AL, PM, and A, respectively. The distributions were shown in the bottom right panels in Figure 3A-C. (C) Correlation matrix showing the tuning similarity (Pearson’s correlation coefficients between SFTF tuning maps) across HVAs (AL, PM and A) and their L2/3 intracortical input pathways. HVAs and their respective input pathways are grouped and shown in the boxes.

**Figure S4.**
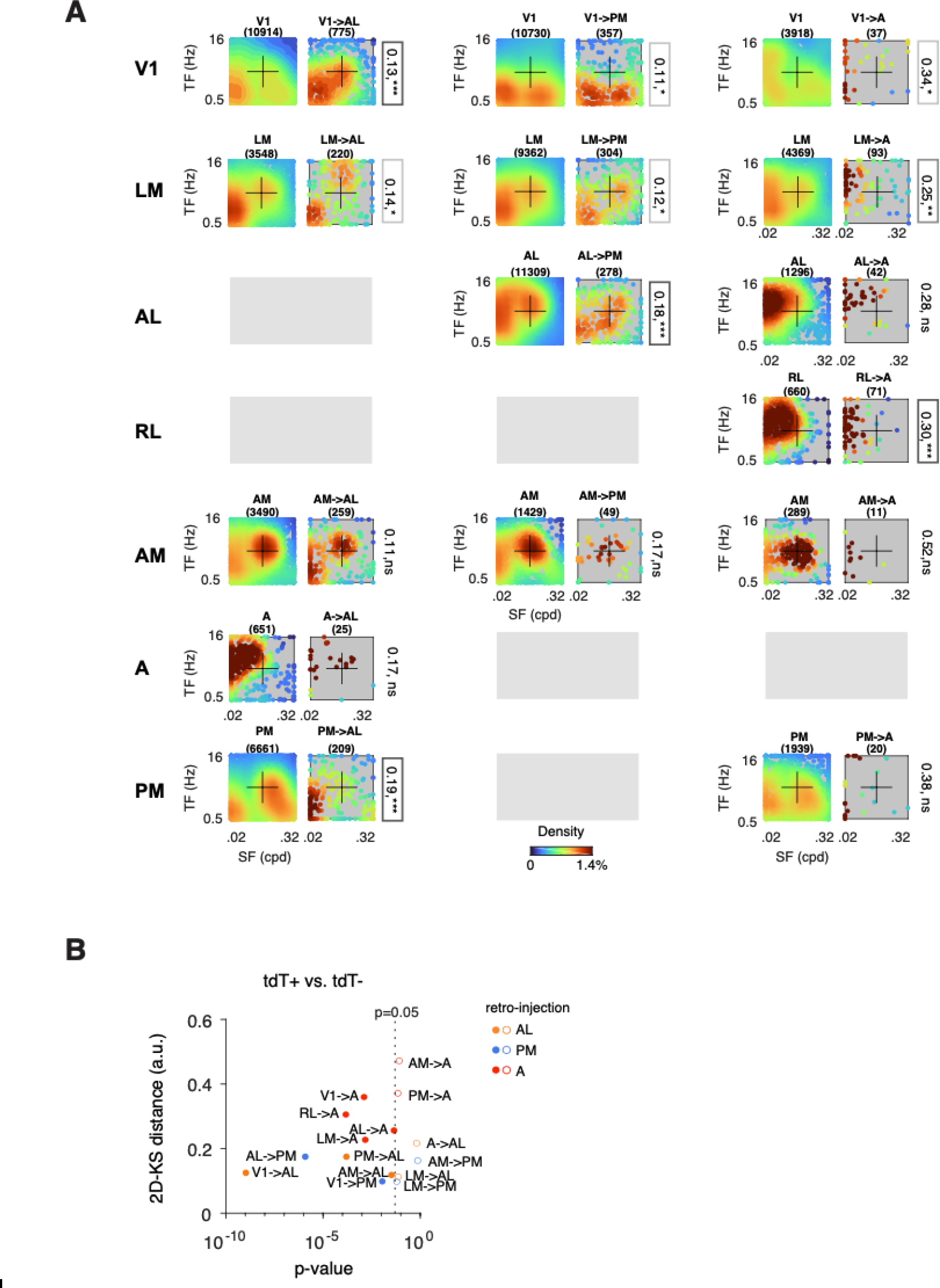
Statistical comparisons of the peak spatiotemporal frequencies between subpopulations of L2/3 intracortical neurons, related to Figure 4. (A) Density scatter plots showing the distribution of peak spatiotemporal frequencies of retrogradely labeled L2/3 projection neurons and non-labeled L2/3 neurons in different areas (by rows) with retrograde virus injections in AL, PM, or A (by columns). Datapoints are color-coded by estimated local density. The density maps are replotted as the contour plots in Figure 3. 2D-KS distances of the distributions of peak spatiotemporal frequencies between tdT- and tdT+ were shown on the right. The significance levels were shown as * and the boxes. p<0.001: ***; p<0.01:**;p<0.05:*;n.s: no significance. (B) Scatter plot showing the p-value and 2D-KS distances between the tdT- and tdT+ neurons in each dataset. Datasets with significant differences (p<0.05, dash line) are shown as solid dots, otherwise circles.

**Figure S5.**
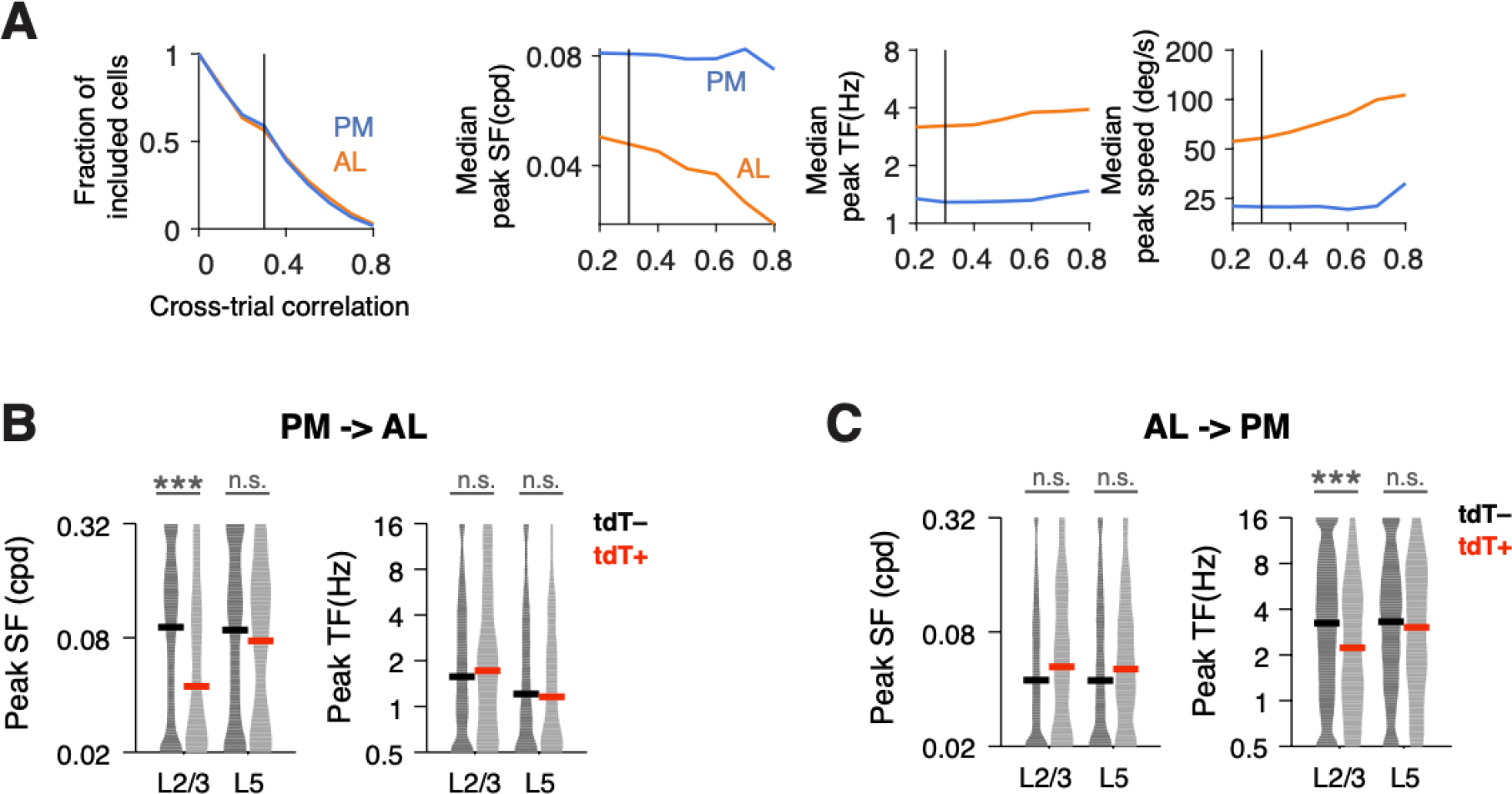
Functional characterization of L5 neurons, related to Figure 5. (A) Effect of the cell selection threshold (minimal cross-trial variability) on the fraction of included neurons, peak spatial frequency, peak temporal frequency, and peak speed. (B) Distribution of peak SF and peak TF of tdT- and AL-projecting tdT+ neurons in L2/3 and L5 of area PM. Horizontal bars: median values. (C) Same as (B), for tdT- and PM-projecting tdT+ neurons in area AL.

**Figure S6.**
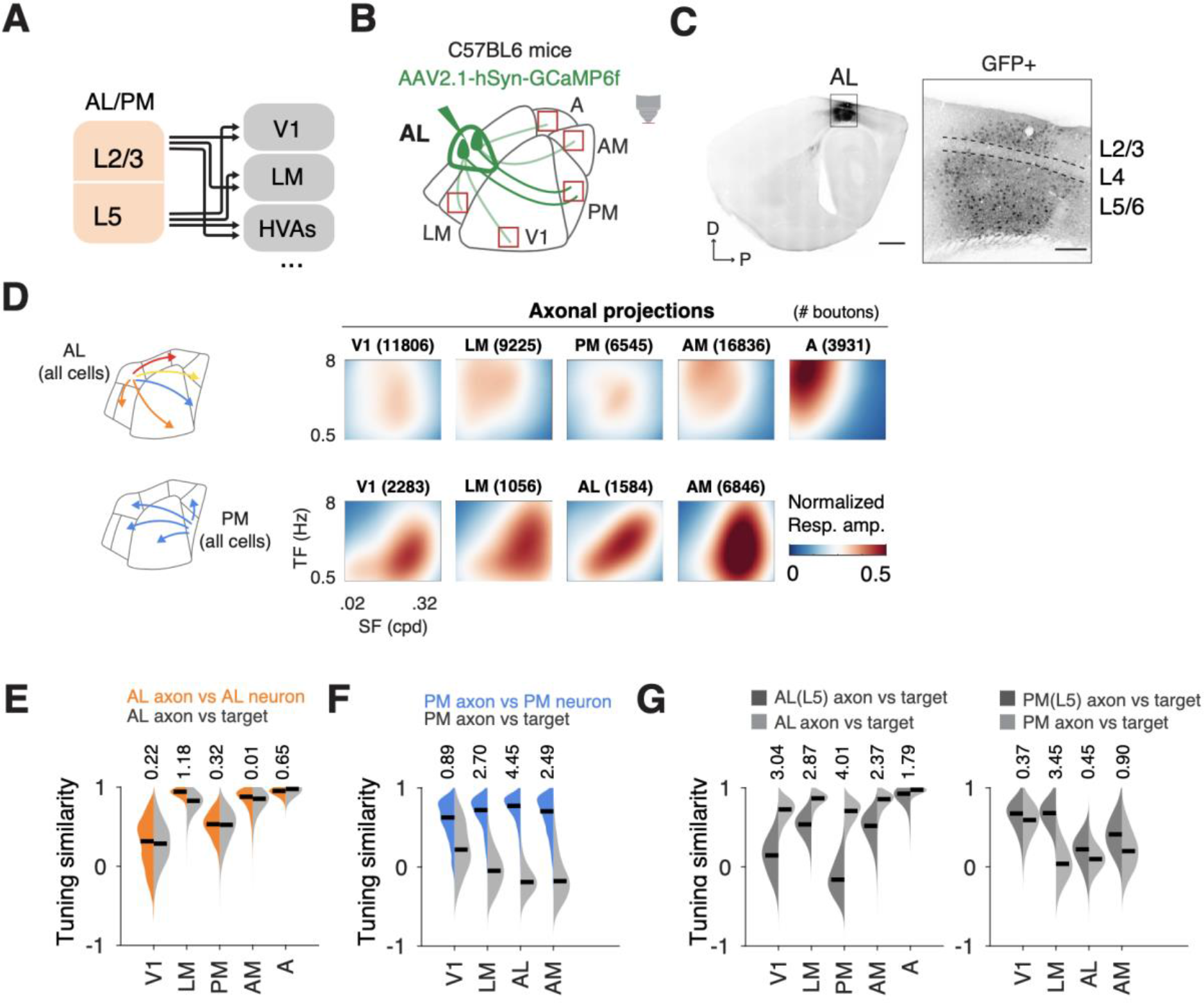
AL, not PM, sends overall target-specific information to distinct cortical areas, related to Figure 6. (B) Schematic showing the convergency of L2/3 and L5 pathways in other cortical areas. (C) Schematic showing the approach to label axons from all projection neurons in area AL by targeted viral injections. Labeled L5 and axonal projections were imaged at across cortical areas using 2P imaging (red boxes). (D) Sagittal slice showing the laminar location and spread of labeled neurons. Scale bar: 1mm. Right: zoom-in at the injection site (black box). Scale bar: 200um. (E) Average spatiotemporal tuning maps of the axonal projections from AL or PM in different cortical areas. (F) Bootstrap distribution of tuning similarity of the AL axons to AL neurons (orange) and to neurons in the target areas (gray). Cohen’s d values are shown on the top of the distributions. (G) Same as (E), for PM axons. (H) Bootstrap distribution of tuning similarity of the AL (or PM) axons, either from L5 neurons (dark gray) or from multi-layer neurons (light gray), to individual target areas. Cohen’s d values are shown on the top of the distributions.

**Figure S7.**
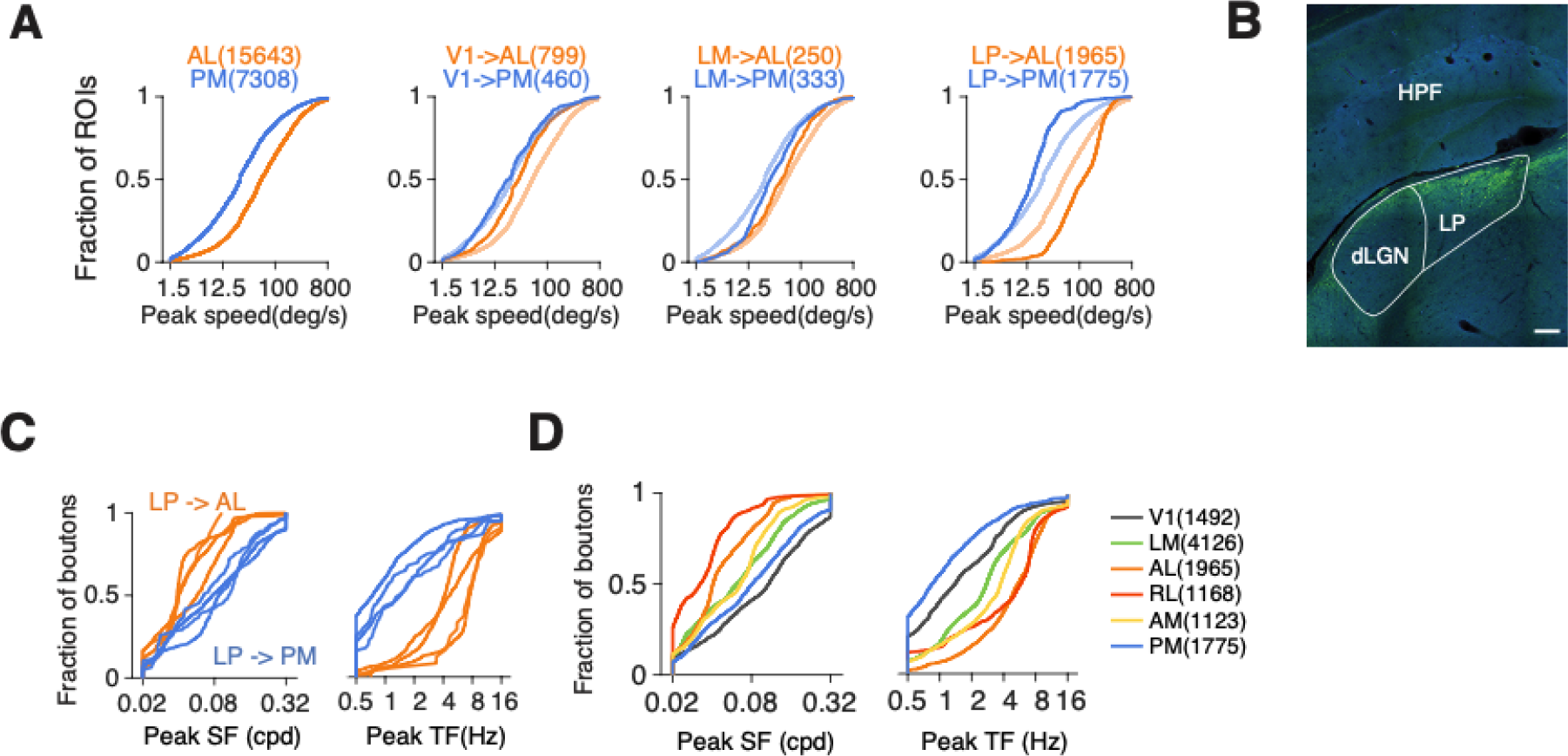
LP conveys specific information to higher visual areas, related to Figure 7. (A) Cumulative distributions of peak speeds of (from left to right) neurons in AL or PM, AL-projecting and PM-projecting neurons in V1, in LM, and LP axons. (B) Example image of GCaMP6 expression after stereotaxic injection in LP. (C) Cumulative distributions of peak spatial and temporal frequencies of LP axons projecting to AL or PM in different mice (four mice for each group). (D) Cumulative distributions of peak spatial and temporal frequencies of LP axons projecting to different cortical areas.

